# *MCM2* mediates post-MI cardioprotection by promoting the pro-angiogenic cardiosome signaling

**DOI:** 10.1101/2024.12.12.628232

**Authors:** Perwez Alam, Wei Huang, Mike Adam, Michelle Nieman, Spencer Stutenroth, Michael Tranter, Yi-Gang Wang, Onur Kanisicak

## Abstract

**Background:** In the past decade, induced cardiac rejuvenation has emerged as a leading approach to repair cardiac injury. Recent studies demonstrate that promoting cell cycle reentry in adult cardiomyocytes (CM) enhances cardiac rejuvenation by influencing paracrine signaling. We previously demonstrated that the inhibition of two cell cycle inhibitors, *Retinoblastoma 1* (Rb1) and *Meis homeobox 2* (Meis2), in the adult CM enhances angiogenesis and cardiac function following ischemic injury, but the underlying mechanisms have yet to be elucidated. The goal of this study is to determine the mechanisms by which inhibition of *Rb1* and *Meis2* promotes cardiac rejuvenation in a mouse model of myocardial infarction.

**Methods:** Myocardial infarction was induced in adult C57/Bl6 mice via permanent LAD occlusion followed by direct injection of either control or *Rb1*+*Meis2* siRNA cocktail to the ischemic myocardium. *MCM2* overexpression done *via* direct myocardial injection of an *MCM2-*plasmid DNA expression cassette. Cardiac function and LV wall motion was assessed via echocardiography, and fibrosis and CM hypertrophy were assessed via histology. RNA-sequencing was performed on isolated adult murine CMs with siRNA-mediated *Rb1*+*Meis2* knockdown to delineate the downstream mechanisms. Further identification of *Rb1, Meis2,* and *MCM2*-dependent mechanisms were done using *in vitro* techniques in isolated CMs and HUVEC cells.

**Results:** We show that siRNA-mediated knockdown of *Rb1* and *Meis2 in vivo* reduces pathological LV remodeling and preserves cardiac structure and function in adult mouse hearts post-MI. RNA-seq analyses revealed *MCM2* as a potential downstream target of *Rb1/Meis2* to enhance protective paracrine signaling in primary adult CMs. Indeed, re-expression of *MCM2,* which is developmentally lost from neonatal to adult CM in the heart, improves cardiac function and LV wall motion while reducing myocyte hypertrophy and fibrotic scar size post-MI. Mechanistically, re-expression of *MCM2* promotes the secretion of pro-angiogenic factors from adult CM, and transfer of conditioned media from *MCM2* expressing CM induced vasculogenesis in HUVEC cells. Proteomic analysis of the *MCM2* interactome confirmed a significant enrichment of angiogenic mediators and suggests an *MCM2*-dependent protein packaging of pro-angiogenic factors in CM-derived small extracellular vesicles (cardiosomes).

**Conclusion:** Re-expression of *MCM2* in adult CM promotes the secretion of pro-angiogenic cardiosomes that induce paracrine revascularization of endothelial cells and mitigates cardiac injury post-MI.

**SUMMARY DIAGRAM:** 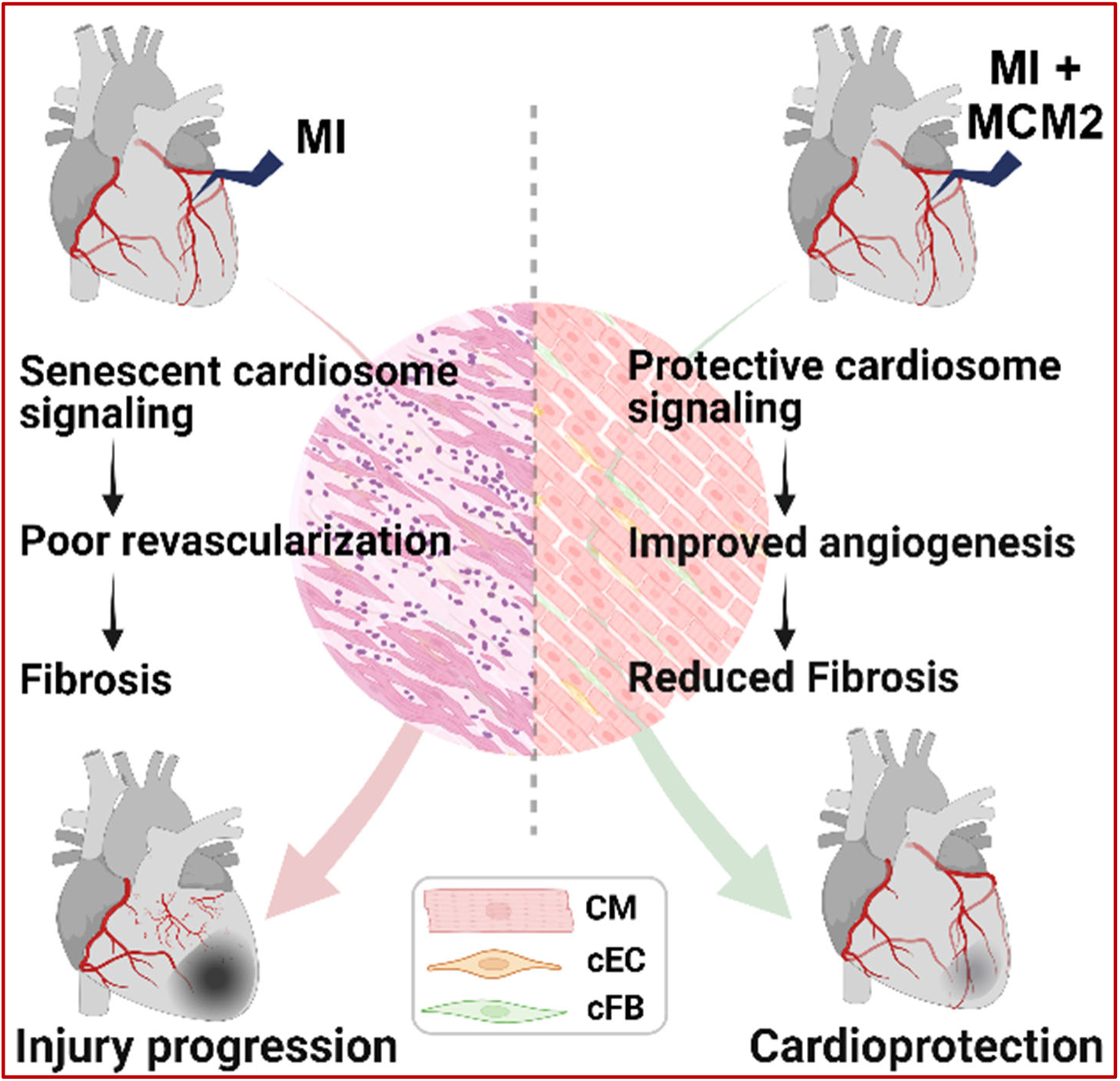

## Introduction

Restricted blood flow is a common pathological factor for ischemic diseases, including myocardial infarction (MI) ^1,2^. MI is a condition caused by the occlusion in coronary vessels, which restricts oxygen, and nutrient supply to the myocardium resulting in cardiomyocyte (CM) death or damage^3,4^. Given that CM are the primary contractile units of the heart, the loss of CM leads to a significant reduction of cardiac function and heart failure^5^. Small animals such as newts and zebrafish have the lifelong ability to regenerate their heart after injury ^6,7^. Similarly, neonatal mouse and to some extent neonatal pig can also repair their hearts after injury ^8,9^. However, unlike zebrafish, newts, neonatal mice or pig, the senescent phenotype of adult CM restricts regenerative capacity of adult mammals’ heart after injury. Therefore, an injury in the adult mammalian heart is considered permanent by current standard of care and leads to adverse cardiac remodeling, presenting an urgent clinical need for therapeutic intervention to either preserve or enhance the number of functional CM following MI.

Induced CM proliferation and cell-based interventions to repopulate CM within the scarred myocardium have been the primary focus of various cardioprotective approaches ^10–14^. However, the current approaches have thus proven inadequate to regenerate the heart with new contracting myocardium to a physiologically, meaningful extent ^15^. Indeed, a common readout of these studies is that the protective effect comes from the protective paracrine signaling that rejuvenate the injured microenvironment, which enhances angiogenesis, reduces fibrosis, and increases CM survival ^16–20^. Studies demonstrate that a major part of the protective effect from stem cell transplantation is derived from their secretome-dependent improvement in angiogenesis, cell viability, and reduced inflammation ^21^. In fact, the administration of stem cell derived secreted factors also improves the cardiac function and reduces infarct size after MI ^22^.

Although repopulating the scar with new functional CM is the ultimate goal to repair the injured myocardium, alternative approaches, such as improving angiogenesis, have evolved as promising approaches to improve cardioprotection ^23,24^. Moreover, therapeutic angiogenesis-based approaches to restore the blood supply to infarcted areas have shown significant improvement in cardiac function and cardioprotection in the adult heart after injury ^25–29^. We have previously reported that reducing CM senescence through siRNA-mediated knockdown of *Retinoblastoma* (*Rb1*) and *Meis homeobox 2* (*Meis2*) improves the cardiac function and reduces the scar size in adult rats after MI through improved angiogenesis ^30^. Our previous work further showed that the activation of the cell cycle improves the pro-angiogenic content of the CM-derived exosomes (cardiosomes). Therefore, enhancing endogenous cardiosome-mediated proangiogenic signaling by manipulating cell cycle-associated genes in the heart represents a novel approach to augment cardioprotection in failing hearts.

Here, we hypothesized that the inhibition of senescence in CM will promote proangiogenic cardiosome signaling and facilitate rejuvenation within the infarcted myocardium. Our results show that the expression of the E2F-dependent cell cycle-associated gene *Minichromosome Maintenance Complex Component 2* (MCM2), which is highly expressed in neonatal CM, is increased following simultaneous inhibition of the *Rb1* and *Meis2* in adult CM ^31^. Reintroduction of *MCM2* expression in the adult heart after MI promotes cardiac rejuvenation and preserved function through increased paracrine secretion of pro-angiogenic cardiosomes. Thus, this work not only supports *MCM2* and CM cell cycle reintroduction as a viable therapeutic target to alleviate pathological cardiac remodeling post-MI, but also highlights the mechanistic importance paracrine cardiosome signaling in promoting angiogenesis.

## Results

### Inhibition of *Rb1* and *Meis2* protects the adult heart after infarction

To confirm the therapeutic significance of *Rb1* and *Meis2* knockdown-mediated cardioprotection, MI was induced in adult (> 12 weeks) mice through permanent LAD ligation followed by intramyocardial injection of *Rb1* and *Meis2*-targeting siRNA (siCocktail) (Figure 1A). Animals receiving siControl post-MI show a MI-dependent increase in CM cell size, whereas treatment with siCocktail blunted this observed increase post-MI (Figure S1). Echocardiographic assessment of cardiac structure and function further demonstrate a preservation of LV ejection fraction along with reduced LV chamber dilation and wall thinning in the siCocktail vs. siControl group post-MI (Figures 1B-E; Figure S2). Immuno-histology analysis further confirms a reduction in infarct size and collagen deposition upon knockdown of *Rb1* and *Meis2* compared to siControl, post-MI (Figures 1F & 1G; Figure S3). Additionally, assessment of vascular:density in the peri-infarct region through αSMA and vWF immunostaining shows a significant increase in the vascular density in siCocktail-injected hearts when compared to siControl after MI (Figures 1H & 1I). Altogether these results corroborate our previous findings, which demonstrate that the inhibition of the *Rb1* and *Meis2* mediate cardiac rejuvenation through promotion of angiogenesis in adult rats post MI ^30^.

**Figure 1:**
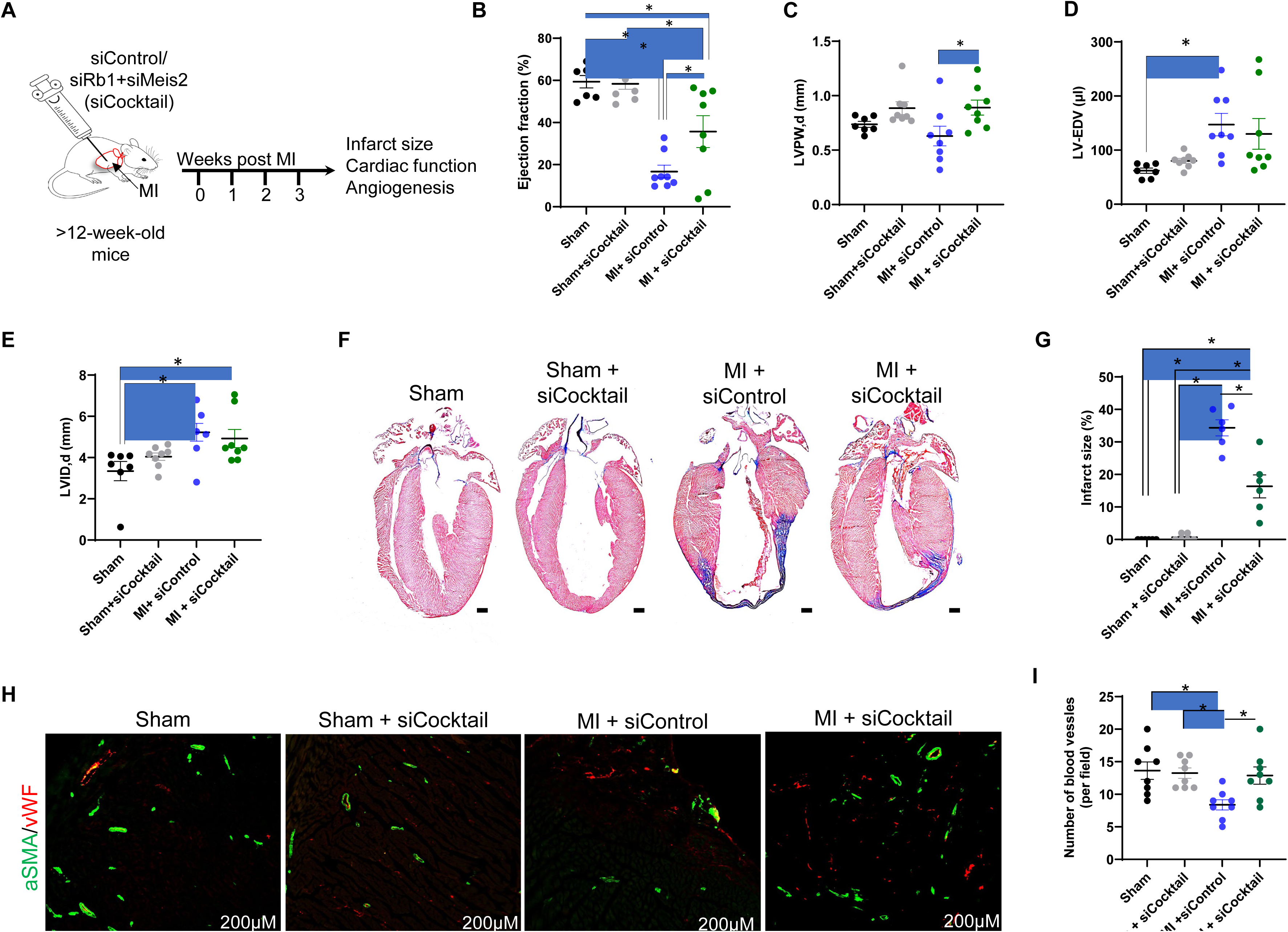
Knocking down Rb1/Meis2 protects cardiac function after MI. (A) Schematics show that knocking down Rb1 and Mesi2 improves cardioprotection. (B) Comprehensive flowchart showing the *in vivo* experimental design, illustrating LAD ligation, intramyocardial injection of siRb1 and siMeis2 or siControl. Animals were observed for 21 days before euthanization and sample collection for molecular and histochemical analysis. Trans-thoracic echocardiography was performed on day 3 and day 21, post- surgery. (C) The dot plot represents the heart-to-body weight ratio among the study groups. (D) Dot plot showing the quantification of ejection fraction among the study groups. (E) Representative Masson’s trichrome-stained images show cardiac remodeling among the study groups. (F) Dot plot representing the quantitative analysis of cardiac remodeling among the groups, which shows reduced infarct size after knocking down of *Rb1*+*Meis2* compared to controls. (G) Vascular density analysis was performed through immuno-histological staining for αSMA (green) and vWF (red). (H) Dot plot showing the quantification of vascular density among the groups. N=6 mice per group. Data represented as mean±SE. * = p-value ≤0.05. # = statically nonsignificant. Scale bars represent the magnification of the corresponding image. Statistical analysis: *T*-test was used to compare the two groups. One-way ANOVA was used to compare multiple groups, and post-hoc analysis (Bonferroni test) was performed to correct the p values from multiple comparisons. *P* value ≤ 0.05 was considered statistically significant. siCocktail = siRb1+ siMeis2.

### *In vitro* inhibition of *Rb1* and *Meis2* activates rejuvenation-associated pathways in adult cardiomyocytes

To determine the downstream mechanisms of *Rb1* and *Meis2* in CM, primary adult murine CM were isolated and transfected with siCocktail followed by RNA-seq analysis at 2- and 7-days post-transfection to respectively analyze the early and late responsive genes (Figure 2A). In addition to assessment of differentially expressed genes (DEG) between siCocktail vs. siControl groups at both time points, we also performed a DEG analysis in the siControl group at 2 (D2) and 7 (D7) days post-transfection (Figure 2B). This comparison not only provides insight into the gene expression response of CMs to the long-term *in vitro* culture, but also allows us to filter out the time-dependent changes observed in the control group and deduce the specific effect of knocking down of *Rb1* and *Meis2* in CM.

**Figure 2:**
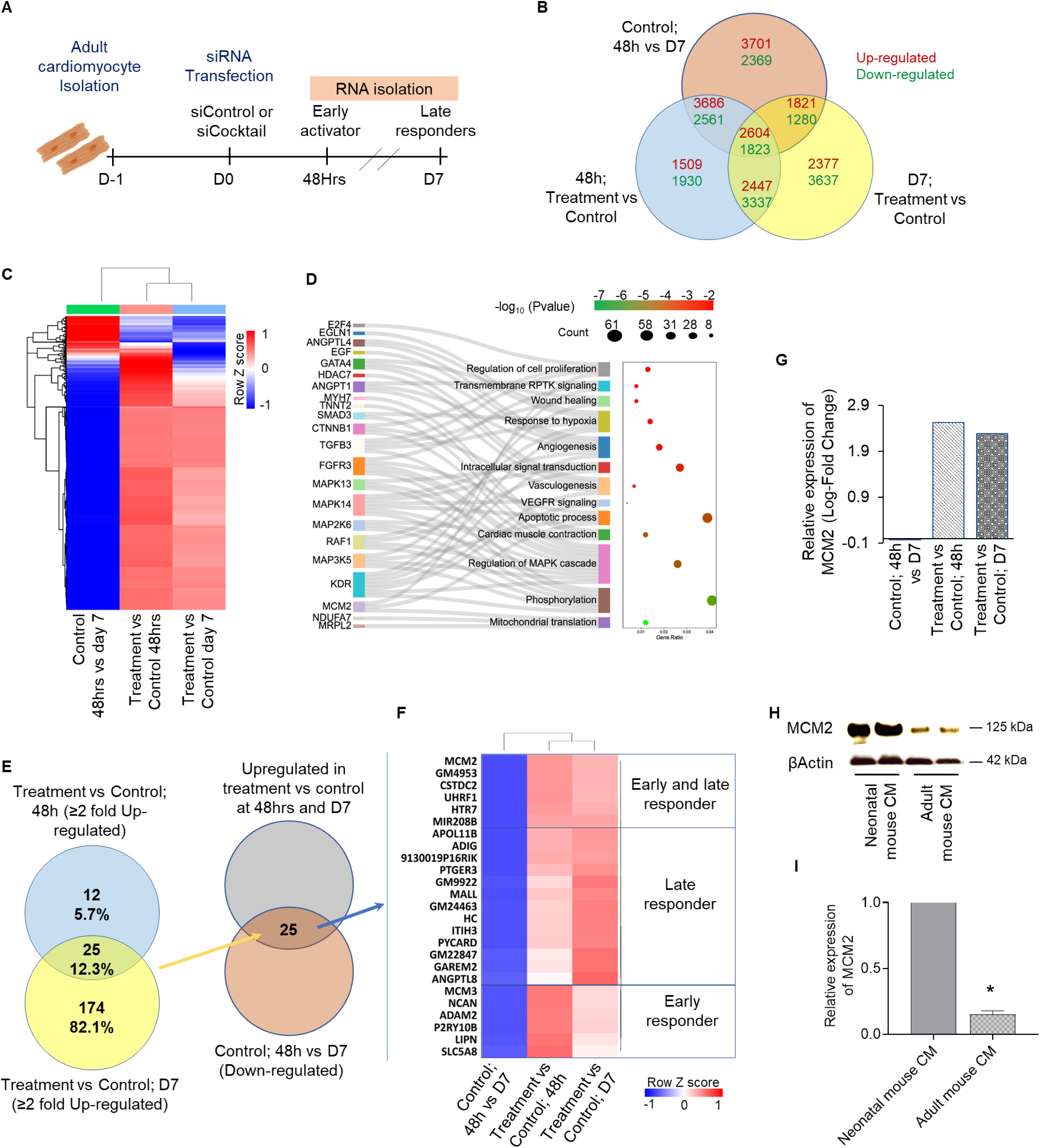
(A) Comprehensive flowchart of adult cardiomyocyte isolation, transfection with siCocktail or siControl, and RNAseq analysis. (B) Bar graph representing the significant knockdown of *Rb1* and *Meis2* after siRNA transfection. (C) Venn diagram represents the comparative analysis approach to analyze the differentially expressed genes among the groups. (D) Heatmap demonstrates differentially expressed genes after knocking down *Rb1* and *Meis2* at early and late time points *versus* control. (E) Sankey and dot plot represents the pathways associated with the up-regulated genes from both early and late-responding groups. (F) Venn diagram showing the 25 genes, commonly up-regulated with ≥2-fold change in siCocktail transfected group (at 48h and D7 after transfection). This analysis further revealed that all 25 genes were downregulated in the control groups at both time points (48h vs D7 post-transfection). (G) Heat map showing the expression of all 25 genes (identified in sub-panel ‘F’) among different study groups. (H) The bar graph shows the expression of *MCM2* at early (48h) and late (D7) time points after knocking down *Rb1* and *Meis2*. (I) The immuno-blot images show reduced expression of MCM2 in adult hearts *versus* neonatal hearts, and the (J) corresponding bar graph shows the quantification of Western blot. N=6 mice per group. Data represented as mean±SE. * = p-value ≤0.05. Statistical analysis: *T*-test was used to compare the two groups. *P* value ≤ 0.05 was considered statistically significant. siCocktail = siRb1+ siMeis2.

RNA sequencing analysis revealed a significant number of genes, which differentially expressed in the siCocktail *vs* siControl groups at D2 and D7 post-transfection (Figure 2C). Heatmap analysis demonstrated a surprisingly high abundance of DEGs whose expression was increased in the siCocktail group at both D2 and D7 (Figure 2C). Pathway enrichment analysis using DAVID functional GO (Biological pathway) analysis revealed a significant enrichment in regeneration associated signaling pathways such as angiogenesis, wound healing, hypoxia response, and cell proliferation (Figure 2D).

To identify the strongest potential targets for further validation studies, these up-regulated genes were further shortlisted based on a cut-off of more than two-fold increased expression after siCocktail treatment at both the early (D2) and late phase (D7), while also showing a natural progression of decreased expression in the control culture over the same time frame (Figure 2E). We identified and further sub- characterized 25 downstream candidate genes into early and late responders to *Rb1* and *Meis2* knockdown based on their expression profile at D2 and D7 post-transfection, with *Minichromosome Maintenance Complex Component 2* (*MCM2*) identified as the strongest early and late responder gene, showing significant upregulation at both D2 and D7 following the knockdown of *Rb1* and *Meis2* (Figures 2F & 2G).

### Re-expression of *MCM2* in the adult heart is cardioprotective post-MI

*MCM2* is a highly conserved protein with 904aa in length and acts as an ATP-dependent DNA helicase as part of the hexameric *MCM2-7* complex ^32^. Loss of *MCM2* expression induces G1/S cell cycle arrest, which may have a role in the senescent state of adult cardiomyocytes ^33^. In support of this notion, we observed a significant decline in *MCM2* expression in adult compared to neonatal CMs (Figures 2H & 2I), which further supports the role of *MCM2* in neonatal CM associated phenotype, and its potential role in promoting cardiac regeneration in the adult heart post-injury.

To investigate the direct role of *MCM2* in cardioprotection, we re-introduced *MCM2* expression in the adult murine heart post-MI via intramyocardial injection of an *MCM2* expression cassette at multiple sites within the infarcted area (Figure 3A). Assessment of CM cell size at three weeks post-MI showed a blunting of MI-induced CM hypertrophy in hearts treated with *MCM2* re-expression (Figures 3B & 3C). Echocardiographic analysis showed that *MCM2* re-expression significantly protects cardiac function after MI when compared to MI control (Figures 3D-G). The observed *MCM2*-dependent improvement in cardiac function was further corroborated by improvement in cardiac output and a strong trend toward reduced ventricular dilation and wall thinning (Figure S4). An in-depth analysis of left ventricular wall function through strain analysis also demonstrates significant improvement in left ventricular wall function with regard to radial displacement, radial strain, radial velocity, and radial strain rate in *MCM2* overexpressing animals post-MI when compared to control (Figure S5). Finally, Masson’s trichrome and Picrosirius red staining both show a significant reduction in fibrotic infarct scar area in mice with cardiac *MCM2* re-expression at three weeks post-MI compared to control (Figures 3H & 3I; Figure S6).

**Figure 3:**
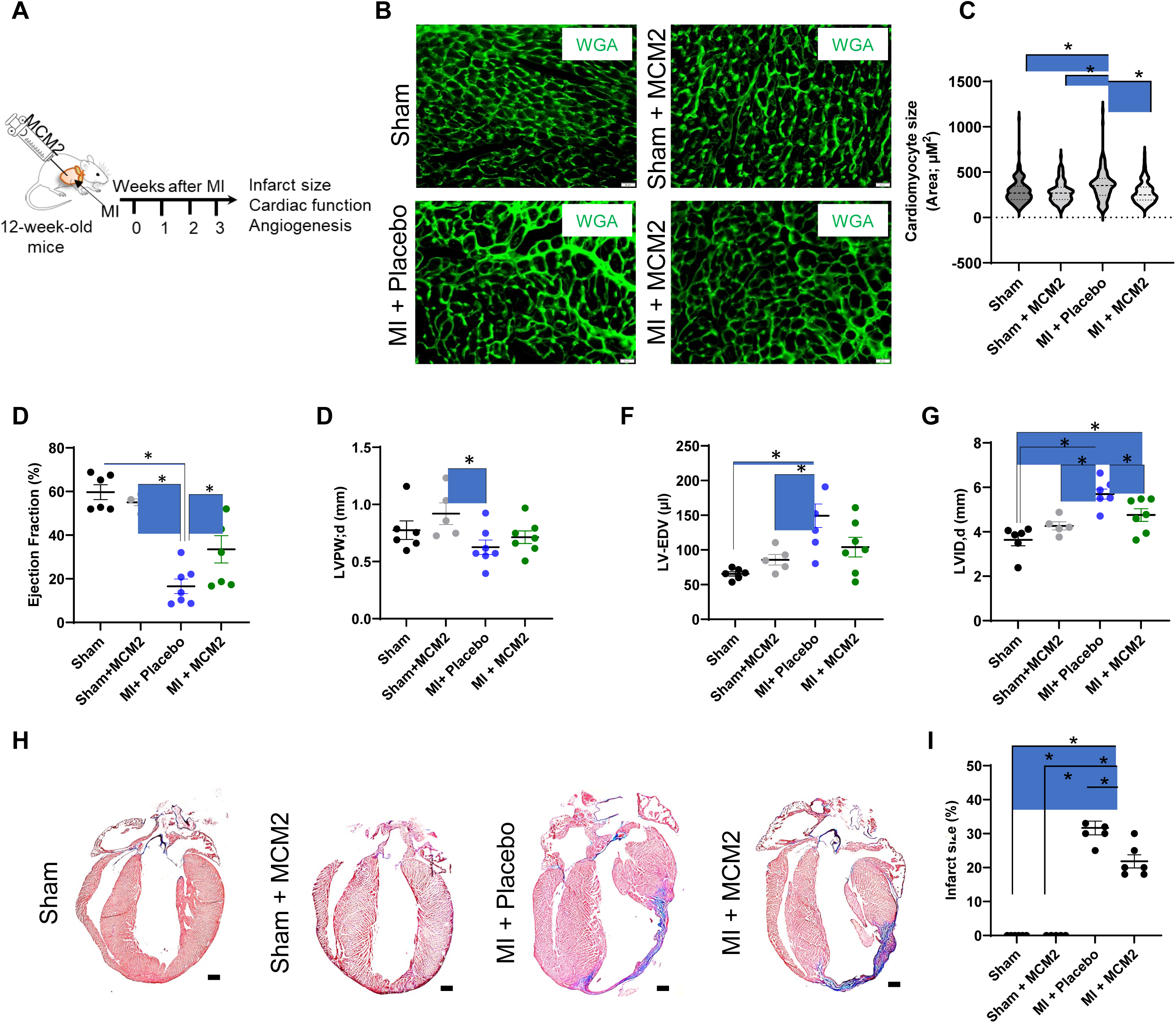
(A) Comprehensive flowchart representing the hypothesis that inhibition of *Rb1* improves cardioprotection through *MCM2*. (B) Schematics showing *in vivo* experimental design, illustrating LAD ligation, intramyocardial injection of *MCM2* overexpressing plasmid, or placebo. Animals were observed for 21 days before euthanization and sample collection for molecular and histochemical analysis. Trans- thoracic echocardiography was performed on day 3 and day 21, post-surgery. (C) Dot plot representation of heart-to-body weight ratio among the study groups. (D) Representative echocardiography images showing the cardiac function in different study groups. Dot plots representing (E) ejection fraction, and (F) cardiac output among the study groups. (G) Representative Masson’s trichrome-stained images show cardiac remodeling among the groups. (H) Dot plot representing the quantitative analysis of Masson’s trichrome stained heart sections in all study groups. N=6 mice per group. Data represented as mean±SE. Scale bars represent the magnification of the corresponding image. Statistical analysis: One-way ANOVA was used to compare multiple groups, and post-hoc analysis (Bonferroni test) was performed to correct the p values from multiple comparisons. *P* value ≤ 0.05 was considered statistically significant. * = p- value ≤0.05. # = statically nonsignificant.

Furthermore, to determine the effect of *MCM2* overexpression on the cell cycle progression, we analyzed the CM proliferation *in vivo* as well as *in vitro*. A reliable identification and analysis of *in vivo* CM proliferation has been challenging. Therefore, we utilized the αMHC-FUCCI mice, a cell cycle visualization reporter mouse, which enables real-time analysis of CMs at different cell cycle states based on the nuclear-specific fluorescent marker, which appears red, yellow, and green for G1, S, and S/G2/M phase, correspondingly^34–36^. Surprisingly, we did not detect an increase in cell cycle activation in CMs post-MI with *MCM2* re-expression compared to the control (Figure S7). Conversely, an *in vitro* analysis using the adult mouse CMs showed a slight but significant increase in *MCM2*-dependent CM proliferation, but this effect was again not detected using neonatal rat ventricular CMs (Figure S8). Overall, these data suggest that induction of CM proliferation is not a significant contributor to *MCM2*-mediated cardioprotection.

### *MCM2* re-expression promotes cardioprotection in the infarcted heart through CM-dependent pro-angiogenic paracrine signaling

Vascular density in the peri-infarct areas was analyzed through fluorescent immunohistochemistry to evaluate the angiogenic potential of *MCM2*. Results showed significantly greater vascular density in heart with re-expression of *MCM2* post-MI (Figures 4A & 4B). However, *MCM2* re-expression had no effect on basal vascular density in mice that received sham surgery. These observations signify the role of *MCM2* in restoring the vascular density and blood supply to infarcted region in failing heart.

**Figure 4:**
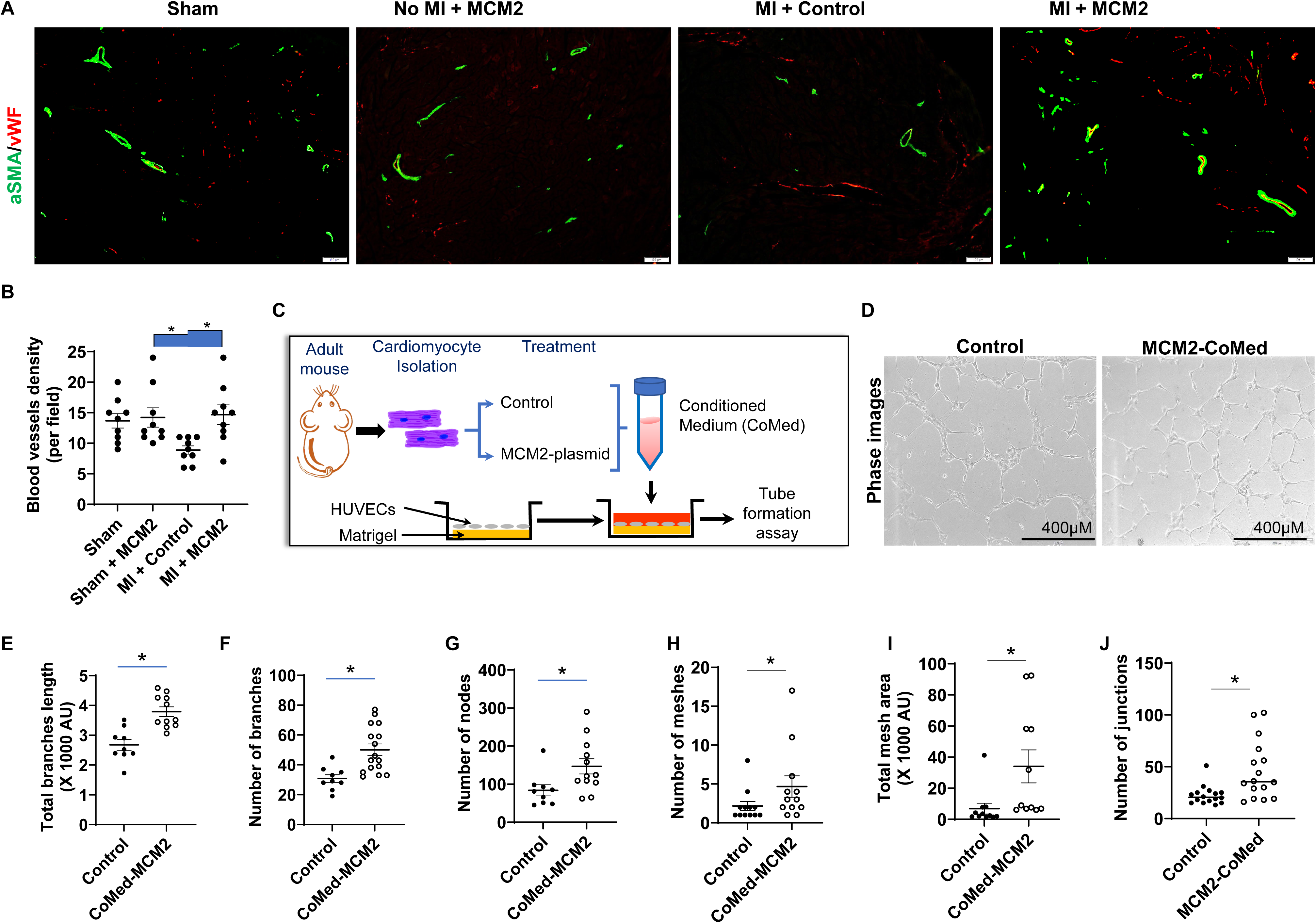
(A) Representative immunohistology images showing vascular density in different study groups. Vascular density was analyzed through αSMA (green), and vWF (red) staining. (B) The dot plot shows the quantification of measured vascular density per field in different study groups. (C) Schematic representation of hypothesis, showing that the paracrine factors from *MCM2* overexpressing adult CMs communicate with endothelial cells and improve angiogenesis. (D) Illustration of tube formation assay, showing ACM transfection and conditioned medium (CM) isolation, which was used to treat HUVECs and the subsequent tube formation assay. (E) Representative bright-field images illustrating endothelial tube formation potential in MCM2-CoMed treated groups *versus* control. (F-H) Dot plots represent the quantification of various parameters of tube formation. Quantification of endothelial tube formation was performed by Image J. Data represented as mean±SE. N = ≥ 10 images per group. Statistical significance was calculated through a *T*-test to compare the data between the groups. However, ANOVA was used to compare the data among the groups, and post hoc analysis (Bonferroni test) was performed to correct the p values from multiple comparisons. *P* value ≤ 0.05 was considered statistically significant. * = p-value ≤0.05. # = statically nonsignificant. Scale bars represent the magnification of the corresponding image. In vivo analysis: N =6 animals per group. In vitro: more than 10 images per group were analyzed for vascular density analysis.

Next, an *in vitro* tube formation assay was done using human umbilical vein endothelial cells (HUVEC) treated with the conditioned medium from primary mouse CM re-expressing *MCM2* (CoMed-MCM2) to determine the angiogenic potential of *MCM2* dependent cardiosomes (Figure 4C). Our results showed a significant increase in angiogenic potential of HUVEC after CoMed-MCM2 treatment *versus* control as assessed through total branch length, number of branches, and number of nodes (Figures 4D-H). We also observed a significant increase in the number of meshes, total mesh area, and number of junctions in the CoMed-MCM2 group (Figure S9A-C), and show that the CM *MCM2*-dependent angiogenic effects are independent of changes in HUVEC proliferation (Figures S9D & S9E). These data demonstrate that re-expression of *MCM2* in CM promotes the secretion of pro-angiogenic factors that promote vascularization of endothelial cells, which improves vascular density and cardioprotection in injured hearts.

### *MCM2* mediates the proangiogenic packaging of CM-derived vesicles through direct protein-protein interactions

To identify the *MCM2* protein interactome within CM to identify potential *MCM2*-dependent mechanisms and signaling pathways of pro-angiogenic paracrine signaling, we utilized the TurboID-based proximity labeling method to identify the *MCM2* interacting proteins (Figure 5A), and identified 243 proteins which showed specific interaction with *MCM2* (Figures 5B & 5C). Pathway enrichment analysis of these proteins revealed a significant association of *MCM2* interacting proteins with angiogenesis-associated pathways (VEGFA-VEGFR2 signaling), in addition to those mediating apoptosis, telomere organization, cell cycle, stress response, and biosynthetic processes (Figure 5D). A cellular component analysis additionally suggested that a significant portion of the *MCM2* interactome are secreted proteins (Figure 5E, Figure S10), strengthening the case for an *MCM2-*dependent secretory pathway as a mediator of post-MI cardioprotection.

**Figure 5:**
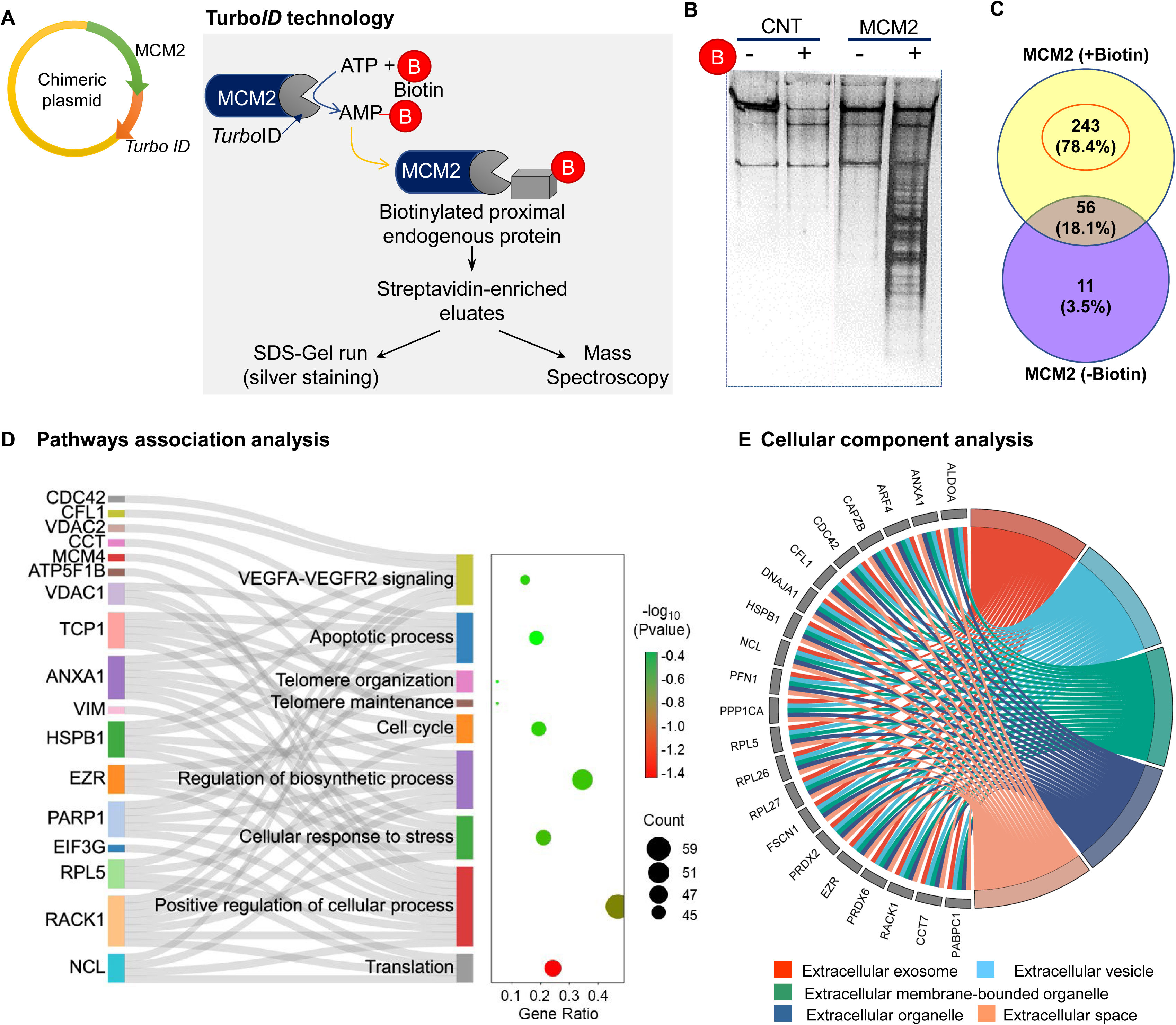
(A) Schematic illustration of protein-protein interaction analysis using TurboID technology. (B) Silver staining image of a 2D SDS-PAGE showing the enrichment of proteins, interacting with MCM2. (C) Venn diagram revealing 243 proteins, interacting with MCM2. (D) Sankey and dot plot shows the biological pathways associated with the proteins that physically interact with MCM2. The size of the bubble corresponds to the number of genes in that component. Whereas, the color of the bubble corresponds to the significance value. (E) The chord plot demonstrates the MCM2 interacting proteins, which also occur in extracellular space and may contribute to paracrine signaling.

Given the high percentage of *MCM2* interacting proteins associated with secreted vesicles, we performed a proteomic analysis of CM-derived small extracellular vesicles (cardiosomes) from *MCM2* expressing and control (GFP) adult CM (Figure 6A). This analysis identified 23 proteins that are uniquely secreted within cardiosomes from *MCM2* expressing compared to control CM (Figure 6B). In accordance with our previous data, 16 of these *MCM2*-dependent cardiosome proteins are also differentially expressed at D2 and D7 after knockdown of *Rb1* and *Meis2* (Figures 2B & 6C). Further protein-protein interaction analysis suggests that 31 of the *MCM2* interacting proteins in CM (identified in Figure 5) directly interact with 13 (out of 23) of the *MCM2*-dependent cardiosome proteins (Figure 6D). Moreover, an *in silico* transcription factor analysis for the *MCM2*-dependent cardiosome proteins predicted that many of them are transcriptionally controlled by transcription factors with known roles in the cardioprotection (NRF1^37^), CM development (CTCF^38^, E2F^31^), hypoxia or stress response (ETS1^39,40^, E2F^30,41^) (Figure 6E). Taken together, these results provides a likely direct mechanism by which *MCM2* re-expression in adult CM drives the pro-angiogenic protein content of cardiosomes post-MI.

**Figure 6:**
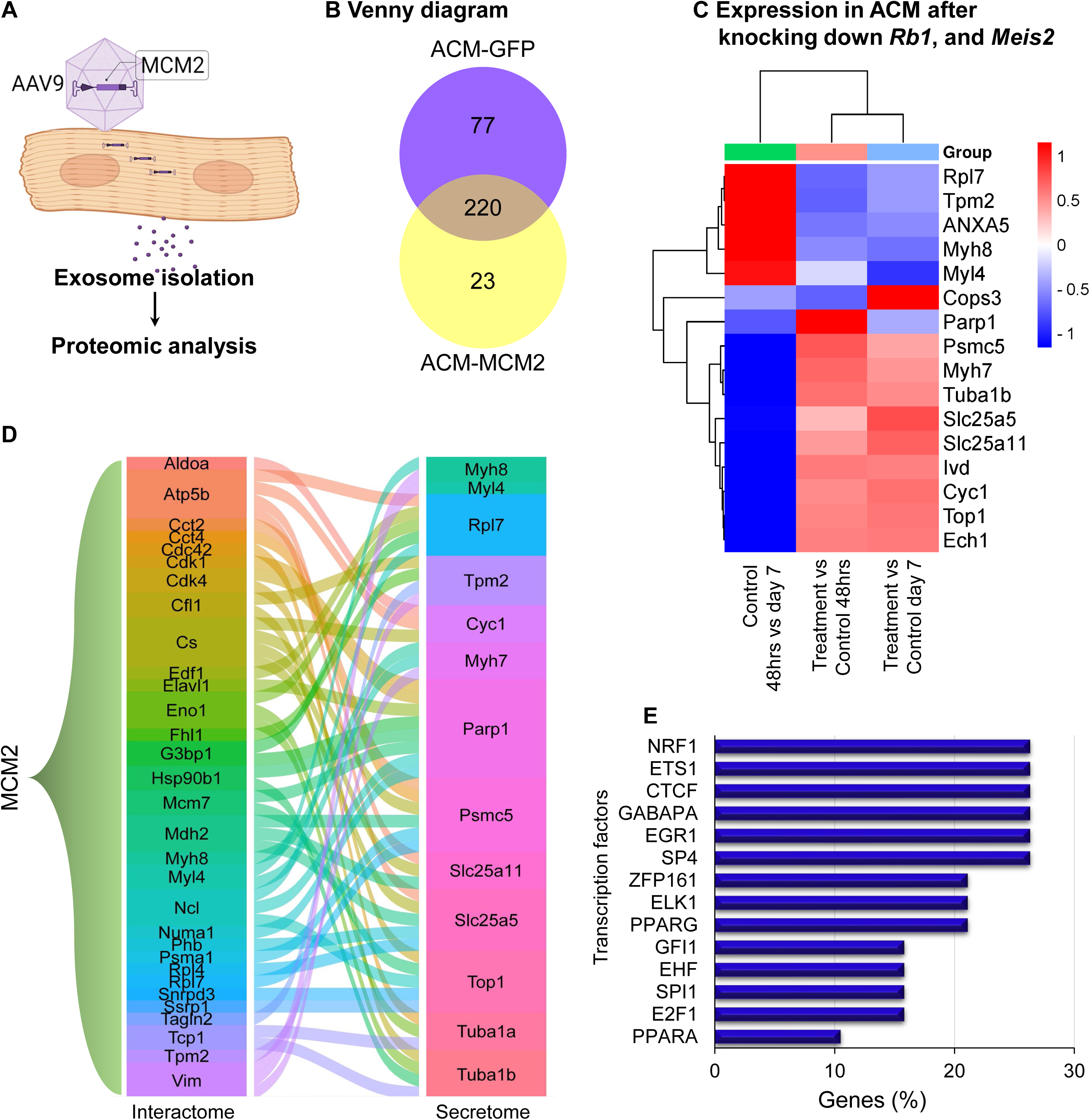
Schematics illustration of *MCM2* overexpression in adult CM using AAV9 viral vector, collection of exosomes from *MCM2* overexpressing CM, and proteomic analysis. The secretory proteins were characterized through Mass spectrometry. (B) Venn diagram revealed 23 unique proteins, which are secreted from the MCM2 overexpressing CM *versus* control. (C) The heatmap shows the expression pattern of the MCM2-induced 23 secretory proteins in previous RNAseq data after knocking down *Rb1* and *Meis2* in adult CM. The expression patterns of all 23 proteins were similar in the RNA seq analysis at early and late time points. (D) *In silico* analysis shows interaction between the 31 MCM2 interacting proteins with 13 proteins, which are secreted from MCM2 overexpressing CM. (E) *In silico* analysis using FunRich3.1.3 shows that many of the 23 proteins, which are specifically secreted from *MCM2* overexpressing CMs are being regulated by hypoxia and stress-associated transcription factors.

## Discussion

Coronary artery occlusion restricts blood supply to the cardiac muscle, limiting oxygen and nutrient delivery to the affected area and causing MI. The senescent nature of cardiomyocytes (CMs) in adult mammals impedes their ability to repair cardiac injury, leading to significant CM death and, over time, resulting in heart failure. Consequently, inducing CM proliferation has become a crucial approach for repairing cardiac damage in adult mammals. However, despite extensive efforts, there has been limited success in replacing cardiac scar tissue with new, functionally active CMs. Research suggests that activating the cell cycle in adult CMs can trigger neonatal-like pathways, enhance intercellular crosstalk through cardiosome-mediated signaling, promote angiogenesis, and reduce adverse remodeling ^42^.

Consistent with these findings, our earlier study demonstrated that inhibiting the senescence-associated genes *Rb1* and *Meis2* protects the adult rat heart post-infarction by activating cardiomyocyte (CM) cell cycles, enhancing angiogenesis, and improving cell survival ^30^. Furthermore, knocking down *Rb1* and *Meis2* in CMs increases the pro-angiogenic content of exosomes, thereby enhancing the angiogenic potential of endothelial cells. The current study elucidates the mechanism by which cell cycle activation in adult CMs modulates extracellular paracrine signaling, specifically CM to endothelial cell (EC) communication, thereby enhancing angiogenesis in the adult heart. This research, therefore, provides molecular insight into how inhibiting senescence in adult CMs enhances protective cardiokine signaling.

Since the previous study was conducted using an adult rat model, we first validated the cardioprotective effect of knocking down *Rb1* and *Meis2* in an adult mouse model of myocardial infarction (MI) before performing mechanistic analyses. Consistent with our earlier findings in the adult rat, we observed that knocking down *Rb1* and *Meis2* in the adult mouse heart enhances cardioprotection after MI. Additionally, we noted a significant increase in vascular density following the knockdown of these genes. These observations support the hypothesis that inhibition of senescence in adult CMs improves angiogenesis and cardioprotection.

Next, we delved into the molecular mechanisms governing the activation of angiogenesis-associated pathways in adult cardiomyocytes (CMs) following the inhibition of *Rb1* and *Meis2*. Employing a comprehensive approach integrating *in vitro* and *in vivo* experiments, our aim was to unravel the intricate signaling networks orchestrating cardiac wound healing and angiogenesis, with a focus on identifying potential therapeutic targets for enhancing cardiac protection post-injury.

Our investigation commenced with the isolation of primary CMs from adult mice, followed by the inhibition of *Rb1* and *Meis2* expression using a cocktail of siRNAs. Building upon our previous study, where we noted cell cycle activation in adult CMs seven days after siRNA-mediated inhibition of *Rb1* and *Meis2*, indicating the heightened responsiveness of adult CMs to external interventions, we conducted data collection at two time points: 24 hours and seven days post-siRNA transfection, to capture both early and late responder genes.

Through RNA-seq analysis at different time points post-transfection, we uncovered significant alterations in gene expression patterns associated with key pathways involved in cardiac repair and regeneration. Notably, our analysis unveiled a robust upregulation of genes implicated in angiogenesis, wound healing, hypoxia response, and cell proliferation following *Rb1* and *Meis2* knockdown, shedding light on the molecular mechanisms underpinning enhanced cardiac rejuvenation. Among the myriad of genes affected by *Rb1* and *Meis2* inhibition, *Minichromosome Maintenance Complex Component 2* (*MCM2*) emerged as a central regulator with potential implications for cardiac rejuvenation. Our data revealed a significant upregulation of *MCM2* at both early and late time points following *Rb1* and *Meis2* knockdown compared to control.

The Rb1/E2F pathway likely acts as the central mechanism inducing *MCM2* expression following si- cocktail treatment. The interaction between Rb1 and E2F factors influences the transcriptional activation of various cell cycle-associated genes, including *MCM2*, by regulating their translocation to the nucleus ^43^. This notion is strongly supported by our previous and current observations, which respectively demonstrate enhanced *E2F* and *MCM2* expression after the knockdown of *Rb1* and *Meis2*. This suggests a potential mechanism through which *Rb1* inhibition may foster cardioprotection by activating *E2F*- mediated pathways and subsequently upregulating *MCM2*, emphasizing the significance of the Rb1/E2F/MCM2 axis in cardiac cellular processes and potential therapeutic interventions for cardiac rejuvenation.

*MCM2* is a highly conserved protein, crucial for DNA unwinding, plays a pivotal role in DNA replication and the cell cycle^32,44^. Inhibiting *MCM2* expression leads to G1/S cell cycle arrest, aligning with the senescent state of adult cardiomyocytes^33^. A study by Rebecca et al. demonstrates elevated expression of both *E2F* and *MCM2* in embryonic and neonatal cardiomyocytes (CMs), while their levels are reduced in adult CMs ^31^. Consistent with these findings, our previous and current results respectively reveal lower *Rb1* expression and increased *MCM2* expression in neonatal hearts compared to adult CMs.

The elevated expression of *MCM2* in neonatal hearts, coupled with its upregulation following the inhibition of *Rb1* and *Meis2* in adult CMs, highlights its critical role in activating neonatal-like pathways, which enhances regenerative and protective capacities in the adult heart. Overall, it suggests the key role of *MCM2* in cardiac protection after injury. This notion is further supported by studies investigating the role of *MCM2* in transcriptional regulation and phosphorylation processes, beyond its established function in DNA replication ^45, 46^.

*MCM2* interacts with the Pol II holoenzyme, affecting the transcriptional activity of various genes. Its transcriptional impact varies; some studies show that *MCM2* blocks RNA Pol II binding at transcription start sites, inhibiting cilia transcription in non-cycling human fibroblasts, while others find that antibody-mediated suppression of *MCM2* inhibits RNA Pol II-mediated transcription in Xenopus oocytes ^47^ ^48^ . Additionally, *MCM2* has been implicated in the phosphorylation of numerous gene sites. Hoi et al. identified numerous phosphosites significantly regulated by *MCM2* overexpression or silencing ^46^. Our TurboID data reveal interactions with four genes (NCL, MAPRE1, RBM14, and PA2G4 or EBP1) where MCM2 may regulates phosphorylation.

These observations suggest that *MCM2* may play a broader role in cellular functions beyond proliferation, contributing to cardiac protection and repair mechanisms. By modulating the expression and activity of *MCM2*, it may be possible to develop therapeutic strategies aimed at rejuvenating the adult heart, improving its ability to repair and regenerate after injury.

Subsequent experiments were conducted to evaluate the cardioprotective effects of *MCM2* in adult mice following myocardial infarction (MI). Our findings from MCM2 overexpression studies revealed significant improvements in cardiac function and reduced scar formation post-MI, underscoring the therapeutic potential of *MCM2* in mitigating cardiac injury. Notably, echocardiographic and histological analyses provided further support for the functional and structural benefits induced by *MCM2* overexpression, affirming its role in promoting cardiac protection. While *in vitro* analyses showed a modest increase in cardiomyocyte proliferation following *MCM2* overexpression, *in vivo* studies using cell cycle visualization reporters did not show significant differences in cardiomyocyte proliferation post-MI. However, we did observe a notable increase in blood vessel density in the peri-infarct region after *MCM2* overexpression. These observations emphasize the intricate nature of cardiac regeneration and underscore the predominant role of non-proliferative mechanisms in mitigating cardiac scarring.

Given the well-established role of paracrine signaling in enhancing angiogenesis in the injured heart, we shifted our focus to elucidating the mechanism of MCM2-dependent pro-angiogenic signaling. To this end, we conducted an in vitro tube formation assay to evaluate the angiogenic potential of factors secreted by MCM2-overexpressing cardiomyocytes (CMs). Consistent with our previous findings using siRb1 and siMeis2, we observed a significant enhancement in the angiogenic potential of HUVECs treated with conditioned media from MCM2-overexpressing adult CMs.

Since the angiogenic role of MCM2 is novel, we aimed to determine the molecular mechanisms underlying the enhanced angiogenic signaling induced by MCM2 overexpression. We focused on identifying proteins that interact with MCM2, employing a proximity-based biotin labeling method using the TurbID technique. Our proteomic analysis revealed 243 proteins specifically interacting with MCM2. Further in silico analysis demonstrated that these interacting proteins are significantly associated with pathways involved in cardioprotection, including angiogenesis, cell cycle regulation, and cellular stress responses. Notably, many of these MCM2-interacting proteins were identified as secretory in our analysis.

We next analyzed exosomes from MCM2-overexpressing cardiomyocytes (CMs) to identify pro-angiogenic factors that could enhance endothelial cell angiogenesis. Our exosome analysis revealed 23 unique proteins secreted by *MCM2*-overexpressing CMs compared to controls. Among these, 16 proteins were differentially expressed following the knockdown of *Rb1* and *Meis2*, as identified through RNA sequencing. *In silico* interaction analysis indicated that thirteen of these proteins interact with MCM2- interacting proteins identified via the TurboID method. Additionally, transcription factor analysis suggested that many of the proteins secreted by *MCM2*-overexpressing CMs are predicted to be regulated by transcription factors involved in cardioprotection, CM development, hypoxia, and stress response ^37,38^ ^30,31,39–41^. While further functional validation is required, these findings offer valuable insights into how MCM2 may modulate intracellular pathways and intercellular signaling to promote angiogenesis. Specifically, they suggest that MCM2 interacts with intracellular partners to enhance pro-angiogenic mechanisms in cardiomyocytes and influence the angiogenic content of exosomes secreted by MCM2- overexpressing CMs.

Overall, our study highlights the pivotal role of *MCM2* in cardiac protection after myocardial infarction (MI). We demonstrated that silencing the senescence-associated genes *Rb1* and *Meis2* enhances angiogenesis and cardiac protection, a finding that is now validated in both adult rat and mouse models. *MCM2* has emerges as a key regulator that activate neonatal-like pathways, and angiogenic signaling in adult heart. Our analyses reveal that *MCM2* interacts with proteins involved in cardioprotection and angiogenesis, enhancing the pro-angiogenic content of cardiosomes. These findings highlight MCM2 as a potential therapeutic target for improve cardiac protection, paving the way for innovative treatments to improve outcomes in heart failure.

## Materials and Methods

### Animal studies

Adult 12-14 week-old C57BL/6 mice, procured from Jackson Laboratory were used in the study. Alternatively, for the *in vivo* cardiomyocyte proliferation analysis αMHC-FUCCI (donated by Mark A Sussman, Ph.D., San Diego State University). All the experiments, and surgical procedures, performed in this study were approved by the Institutional Animal Care and Use Committee at the University of Cincinnati.

### Myocardial infarction

Myocardial infarction was created by permanent ligation of left anterior descending coronary artery (LAD) in adult (12-14 weeks old) mice, as mentioned before^49^. Briefly, animals were anesthetized by isoflurane gas inhalation. Subsequently, tracheal intubation was performed, and mice were ventilated by using rodent ventilator (Minivent, Type 845, Hugo Sachs Elektronik, March-Hugstetten, Germany). A left sided thoracotomy was performed to expose the heart, and LAD was ligated using the suture to create the MI, as per our standardized protocol^50–52^. Immediately after LAD ligation, siControl or siCocktail, prepared in the Invivofectamine 3.0 reagent (Invitrogen) as per the manufacturer’s protocol, were intramyocardially injected at the multiple sites in the area under infarction. After injection, the chest was closed and animals were followed for 21 days post-surgery, before termination of the experiment.

### Cardiac function analysis

In vivo cardiac function was analyzed through transthoracic echocardiography using Vivo 2100 Imaging digital ultrasound system (VisualSonics), as previously discussed ^50,52^. The animals were anesthetized by isoflurane inhalation. Animals were placed in supine position under 2-3% isoflurane at a heart rate of ∼ 400 beats/min, throughout the procedure. The two-dimensional imaging across the cardiac anterior and posterior wall of left ventricle was performed for B and M mode using a 22–55 MHz (MS550D) transducer. The images were recorded at parasternal cardiac long axis view, at the level of mid-papillary muscles. The cardiac function parameters, such as ejection fraction (EF), fractional shortening (FS), cardiac output (CO), and wall parameters were analyzed from at least 3 consecutive cardiac cycles. Additionally, we performed VivoStrain tool to analyze detailed wall function as mentioned previously ^53^.

### Histological examination and immunohistochemistry

On day 21 post surgery, animals were euthanized, and hearts were collected for histological analysis. Prior to isolation, hearts were perfused with 1M KCl to arrest them in diastolic position and fixed by four hours incubation at 4°C in 4% paraformaldehyde (PFA), followed by immersion in 30% sucrose until the heart sinks. Fixed hearts, then embedded into OCT compound (Tissue Tek), and placed into -80C freezer following the standard protocol ^54^. The hearts were cryo-sectioned at 10µm thickness, for subsequent histological analysis. Masson’s trichrome staining was performed by the histology core at Cincinnati Children’s Hospital and medical center (CCHMC). Picrosirius red staining was performed using the Picro- Sirius Red Solution (Abcam), following the manufacturer’s protocol.

For immune-histological analysis, tissue sections were blocked for 1 hour at room temperature in CAS-Block (Thermo Fisher Scientific, Waltham, MA). Subsequently, the primary antibodies were diluted 1:200 in CAS-Block, and the heart sections were individually incubated overnight with corresponding antibodies (antibody details are provided in supplementary table 1).

### Adult cardiomyocyte isolation and culture, and transfection

Mouse adult CMs were isolated from >12weeks old mice using the modified Langendorff procedure^55^. Briefly, Adult ice were anesthetized by intraperitoneal pentobarbital injection. Subsequently, chest was open to isolate the heart and perfused with the myocyte buffer to clear any blood. Next the heart was attached to the Langendorff apparatus and perfused with myocyte buffer. After 2-3 min, myocyte buffer was replaced with enzyme solution, and the heart was further perfused for another 10-15min to ensure the appropriate digestion. Next, the heart was taken down from the Langendorff apparatus, and remaining procedure was performed in laminar flow hood in stop solution. The upper atrial portion was chopped out, and the ventricular part was minced with fine forceps to get the single cell suspension. The cell suspension was passed through 60uM cell strainer, followed by the centrifugation at 300rpm for 3min to remove the non-myocyte contamination. The isolated single cell adult CM were plated into Laminin coated plate in myocyte culture media.

Next day after isolation, the cells were transfected with equimolar concentration (100nM) of specific siRNAs against *Rb1* and *Meis2* (siCocktail) using the RNAiMax (Invitrogen) transfection reagents following manufacturer’s protocol. Cells transfected with scrambled siRNA served as control. CMs were followed for 48hrs, or 7 days before RNA isolation and subsequent RNAeq analysis (Figure 2A).

### RNA sequencing

Total RNA isolation was isolated using RNA easy kit (Qiagen) as per the manufacturer’s protocol. RNA sequencing was performed by the RNA Sequencing Core at Cincinnati Children’s Hospital and Medical Center (CCHMC). Sequence read mapping and differential gene analysis were performed by Bioinformatics core at CCHMC. The statistical significance threshold for differential expression analysis was defined as an FDR-corrected P-value less than or equal to 0.05 and a fold-change greater than or equal to 2. All RNA-seq data is available in the NCBI Gene Expression Omnibus (https://www.ncbi.nlm.nih.gov/geo) (GSE 275673).

### Real time expression analysis

Transcriptomic expression analysis of target genes was performed by RT-PCR. Total RNA isolation was performed using the RNA easy micro kit (Qiagen) as per the protocol, and cDNA strands were generated using SuperScript™ III Reverse Transcriptase (Invitrogen) kit following the manufacturer’s protocol. RT-PCR reactions were performed by using SYBR Green PCR Master Mix (Applied Biosystems) in CFX Connect Real-Time PCR Detection System (Bio-Rad, Hercules, CA). The quantitative gene expression was analysis through ΔΔCt method, using β-actin or GAPDH as reference.

The specific primers to amplify Rb1 (forward: AATACACTCTGTGCACGCCT; reverse: TGAGCCAGGAGTCTGGTGTC), Meis2 (forward: GCTCCTGGGCACTGATGAAA; reverse: GCTGGGCCTTTTGAAGGTGA), MCM2 (*forward*: ATCCACCACCGCTTCAAGAAC; *reverse*: TACCACCAAACTCTCACGGTT), Actb (forward: GTGACGTTGACATCCGTAAAGA; reverse: GCCGGACTCATCGTACTCC), and GAPDH (forward: GTATGACTCCACTCACGGCAAA; reverse: GGTCTCGCTCCTGGAAGATG) were used to specifically analyze their expression.

### In vitro cardiomyocyte proliferation assay

Adult mouse CMs isolation was performed as mentioned before. The neonatal rat ventricular CM (NRVCM) were isolated from 1-3 days old rats, following the standard protocol ^56^. To induce the MCM2 overexpression these cells were transfected with *MCM2* overexpressing plasmid (Addgene# 54164), using Lipofectamine 3000 reagent (Invitrogen). Cells were maintained for 7 days in cultured before fixing them with 4% PFA, and subsequent immune-fluorescent assays. Adult CMs media was supplemented with 5 μmol/L of EdU (a nucleoside analogue of thymidine). The cell cycle progression in adult CMs by analyzing the DNA synthesis was performed through EdU staining using Click-iT EdU labeling kit (Invitrogen, Carlsbad, CA) according to the manufacturer’s protocol. Whereas in NRVCT, the cell cycle progression was analyzed through synthesis phase marker KI67, as per our standard immunofluorescent staining protocol^30^. The CMs were stained through cardiac specific marker TnI, whereas nuclei were visualized by DAPI staining. The double positive CMs for TnI, and EdU/KI67 were only considered for analysis. All the experiments were performed in triplicate.

### In vitro endothelial cell proliferation assay

To analyze the endothelial cell proliferation in the presence of conditioned media from MCM2 overexpressing CMs (MCM2-CoMed), HUVEC were cultured on in a collagen coated plate. The MCM2- CoMed was collected from the adult CMs transfected with MCM2 overexpressing plasmid. Prior to the MCM2-CoMed treatment, HUVEC were kept in basic media for at least 12 hours to avoid the interference of growth factors on proliferation, which are present in the endothelial cell culture media. As mentioned for NRVCM, the endothelial cell proliferation was also analyzed through KI67 assay. The endothelial cells were specifically stained by Vimentin, and the double positive cells for Vimentin, and KI67 were analyzed to calculate the proliferation. All the experiments were performed in triplicate.

### *In vitro* tube formation Assay

To analyze the pro-angiogenic potential of cardiosomes from *MCM2* overexpressing cells, an *in vitro* tube formation assay was performed by using the HUVEC^30^. HUVEC cells were cultured on Matrigel (Geltrex Reduced Growth Factor Basement Membrane Matrix, Invitrogen, Carlsbad, CA) coated 48 well plates with a seeding density of 42,000 cells/cm^2^. The study was performed in two groups, one group was designated as control, and treated with the conditioned medium from CMs transfected with GFP, whereas the second group was treated with MCM2-CoMed. Cells were cultured in CO2 incubator at 37°C for eight hours before imaging and tub formation analysis. Images were analyzed through Wimasis (2017) image analysis tool (WimTube: Tube Formation Assay Image Analysis Solution, Release 4.0; Onimagin Technologies, Córdoba, Spain). All the experiments were performed three times in triplicate.

### TurboID based proximity labelling and MCM2 interacting protein identification

We utilized TurboID biotin ligase method to identify the proteins interacting with MCM2. TurboID is an engineered biotin ligase that uses ATP to convert biotin into biotin–AMP, a reactive intermediate that covalently labels interacting proteins^57^ (Fig. 5A). Briefly, the full-length cDNA of mouse *MCM2* was amplified from mEmerald-MCM2-N-22 (Addgene ID: 54164) using forward primer: *TAAGCAGAATT CATGCAAGCGGGCCCG*, and reverse primer: *TGCTTAGGATCCGAACTGCTGCAGGATCATTTTC*.

The amplified MCM2 cDNA was cloned into the MCS-BioID2-HA vector (Addgene ID: 74224) using *EcoR1*and *BamH1* restriction sites. Next, AD-293 cells were transfected with the chimeric *MCM2*- BioID2-HA plasmid. After 72 hours of transfection, cells were treated with biotin for 24 hours. Subsequently, protein isolation was performed, and biotinylated proteins were purified using Pierce™ Streptavidin Magnetic Beads (Thermo scientific). Next, the purified proteins were separated on SDS-PAGE, visualized through silver-staining, and unique protein bands were identified through Mass Spectrometry (Fig. 5B). The Mass Spectrometry was performed by the Proteomic core (Biochemistry, proteomics, and biological mass spectrometry) at University of Cincinnati, Cincinnati, USA.

### Exosome analysis

Towards this end, we used AAV9-viral vector-mediated overexpression of *MCM2* in adult mouse CM. First, the AAV9-TNT-*MCM2* vector was created by amplifying MCM2 cDNA from mEmerald-MCM2- N-22 (Addgene ID: 54164) using forward primer: *TAAGCAggatccaccaATGCAAGCGGGCCCGGCC*, and reverse primer: *TGCTTAaagcttGAATTCTCAGAACTGCTGCAG*. The amplified *MCM2* cDNA was then cloned into pAAV-CAG-GFP vector (Addgene ID:37825) by replacing GFP coding sequences using *BamHI*, and *HindIII* restriction sites. Next, to produce the virus, we transfected the HEK293 cells with the pAAV-CAG-*MCM2* chimeric vector along with the pAdDeltaF6 (helper) and AAV2/9n (AAV packaging) plasmids, according to the standard protocol. The purified AAV9-CAG-*MCM2* virus was used to transduce adult CM (Fig. 6A). The AAV9-CAG-GFP virus served as control. On day three after transduction, CM media was replaced with serum-free media. After 48 hours, the conditioned medium was collected, and cardiosomes isolation was performed through ultracentrifugation method following standard protocol^58^. Subsequently, protein isolation was performed from the cardiosomes, separated on SDS-PAGE, and visualized through silver staining using Thermo Scientific™ Pierce™ Silver Stain Kit (Thermo), following the manufacturer’s protocol. Further, proteins from cardiosomes were identified through nano-ESI–MS/MS analysis^59,60^. The Mass Spectrometry was performed by the Proteomic core (Biochemistry, proteomics, and biological mass spectrometry) at University of Cincinnati, Cincinnati, USA.

### Statistical analysis

Data are presented as mean ± standard error mean. The comparative analysis and statistical significance between two groups was calculated by *t*-test. Whereas one-way ANOVA was used to compare multiple groups, and post-hoc analysis (Tukey’s test) was performed to correct the *p* values from multiple comparisons. The p value ≤0.05 was considered as statistically significant.

## Non-standard Abbreviations and Acronym

CMs: Cardiomyocytes
LAD: Left anterior descending (artery)
siCocktail: A combination of siRNAs targeting Rb1 and Meis2
vWF: von Willebrand factor
CO: Cardiac Output.
Co-med: Conditioned Media. EF: Ejection Fraction.
ESI-MS/MS: Electrospray Ionization Tandem Mass Spectrometry.
FS: Fractional Shortening.
MCS: Multiple Cloning Site. MS: Mass Spectrometry.
NRVCMs: Neonatal Rat Ventricular Cardiomyocytes.
OCT: Optimal Cutting Temperature compound.
PFA: Paraformaldehyde.
RT-PCR: Reverse Transcription Polymerase Chain Reaction.
SDS-PAGE: Sodium Dodecyl Sulfate Polyacrylamide Gel Electrophoresis.
siCocktail: A combination of specific siRNAs against Rb1 and Meis2.
siControl: Control small interfering RNA.
TnI: Troponin I
TurboID: Biotin ligase enzyme used for proximity labeling.

---

| **Table 1** | **Antibody** | **Catalog #** | **Company** |
| --- | --- | --- | --- |
| 1 | Rabbit anti-β-actin | A1978 | Sigma-Aldrich |
| 2 | Mouse Anti-αSMA | A5228 | Sigma-Aldrich |
| 3 | Rabbit anti-GAPDH | G9545 | Sigma-Aldrich |
| 4 | Rabbit anti-aurora B | A5102 | Sigma-Aldrich |
| 5 | Mouse anti- cardiac troponin-T | MA5-12960 | Thermo Scientific |
| 6 | Rabbit-anti-Rb1 | 10048-2-Ig | Proteintech |
| 7 | Rabbit anti-cardiac troponin-I | sc-15368 | Santa Cruz Biotechnology |
| 8 | Goat-anti-Meis1/2 | Sc-10599 | Santa Cruz Biotechnology |
| 9 | Rabbit anti-VEGF | sc-152 | Santa Cruz Biotechnology |
| 10 | Rabbit-anti-Bax | 2772S | Cell Signaling Technology |
| 11 | Rabbit anti-β-catenin | 9562S | Cell Signaling Technology |
| 12 | Rabbit anti-p16 | ab51243 | Abcam |
| 13 | Mouse-anti-Histone H3 | Ab6002 | Abcam |
| 14 | Mouse anti-Ki67 | 550609 | BD Pharmingen |
| 15 | ECL anti-mouse-HRP | NA931V | GE Healthcare |
| 16 | ECL anti-rabbit-HRP | NA9340V | GE Healthcare |
| 17 | Goat anti-mouse Alexa fluor 488 | A11029 | Life Technologies |
| 18 | Goat anti-mouse Alexa fluor 594 | A11005 | Life Technologies |
| 19 | Goat anti-rabbit Alexa fluor 488 | A11008 | Life Technologies |
| 20 | Goat anti-rabbit Alexa fluor 594 | A11037 | Life Technologies |

**Supplementary** Figure 1: (A) Representative immunohistology images for Wheat germ agglutinin (green) showing the adult CM cell size in sham, no MI + siCocktail, MI+ siCocktail, and MI + siControl. Nuclei were visualized through DAPI (blue) staining. (B) Graph representing the quantitative analysis of CM size in different study groups. N=6 mice per group. Data represented as mean±SE. * = p-value ≤0.05. # = statically nonsignificant. Scale bars represent the magnification of the corresponding image. Statistical analysis: One-way ANOVA was used to compare multiple groups, and post-hoc analysis (Bonferroni test) was performed to correct the p values from multiple comparisons. *P* value ≤ 0.05 was considered statistically significant. siCocktail = siRb1+ siMeis2.

**Supplementary** Figure 2: (A) Representative echocardiography images showing the cardiac wall function in different study groups. (B-G) Dot plots represent the quantitative analysis of different parameters among the study groups. N=6 mice per group. Data represented as mean±SE. * = p-value ≤0.05. # = statically nonsignificant. Statistical analysis: One-way ANOVA was used to compare multiple groups, and post-hoc analysis (Bonferroni test) was performed to correct the p values from multiple comparisons. *P* value ≤ 0.05 was considered statistically significant. siCocktail = siRb1+ siMeis2.

**Supplementary** Figure 3: (A) Representative Picrosirius red-stained images showing scars in different study groups. (B) Dot plot representing the quantitative analysis of collagen deposition in different study groups. N=6 mice per group. Data represented as mean±SE. * = p-value ≤0.05. Scale bars represent the magnification of the corresponding image. Statistical analysis: One-way ANOVA was used to compare multiple groups, and post-hoc analysis (Bonferroni test) was performed to correct the p values from multiple comparisons. P value ≤ 0.05 was considered statistically significant. siCocktail = siRb1+ siMeis2.

**Supplementary** Figure 4: (A) Representative immunohistology images for Wheat germ agglutinin (green) showing the adult CM cell size in sham, no MI + MCM2, MI+ MCM2, and MI + Control. (B) Graph representing the quantitative analysis of CM size in different study groups. N=6 mice per group. Data represented as mean±SE. Scale bars represent the magnification of the corresponding image. Statistical analysis: One-way ANOVA was used to compare multiple groups, and post-hoc analysis (Bonferroni test) was performed to correct the p values from multiple comparisons. *P* value ≤ 0.05 was considered statistically significant. * = p-value ≤0.05. # = statically nonsignificant.

**Supplementary** Figure 5: (A-E) Representative dot plots showing the quantification of different cardiac function parameters including the wall dimension, and stroke volume. N=6 mice per group. Data represented as mean±SE. Statistical analysis: One-way ANOVA was used to compare multiple groups, and post-hoc analysis (Bonferroni test) was performed to correct the p values from multiple comparisons. *P* value ≤ 0.05 was considered statistically significant. * = p-value ≤0.05. # = statically nonsignificant.

**Supplementary** Figure 6: The representative images showing the analysis of cardiac wall displacement (A-C) and strain (E-G) among the study groups. The graphs show the quantification data for various strain analysis parameters such as displacement (D), strain (H) velocity (I), and strain rate (J) among the groups at day 21 post-surgery. N≥5 mice per group. * = significant versus No MI + MCM2; # = non-significant versus No MI + MCM2; $ = significant versus MI + Placebo; ‡ = non-significant versus No MI + MCM2.

**Supplementary** Figure 7: (A) Representative Picrosirius red-stained images showing scars in different study groups. (B) Dot plot representing the quantitative analysis of collagen deposition in different study groups. N=6 mice per group. Data represented as mean±SE. * = p-value ≤0.05. Scale bars represent the magnification of the corresponding image. Statistical analysis: One-way ANOVA was used to compare multiple groups, and post-hoc analysis (Bonferroni test) was performed to correct the p values from multiple comparisons. P value ≤ 0.05 was considered statistically significant.

**Supplementary** Figure 8: (A) Representative histology images from αMHC-FUCCI animals showing the CMs in advanced stages of the cell cycle (green nuclei) and non-proliferating CMs (red or yellow nuclei) in *MCM2* overexpressing animals *versus* control, after MI. Nuclei were visualized through DAPI (blue) staining. (B) The bar graph represents the quantitative analysis of CM proliferation in different study groups. Data represented as mean±SE. Scale bars represent the magnification of the corresponding image. Statistical analysis: *T*-test for comparing two groups. *P* value ≤ 0.05 was considered statistically significant. # = p-value >0.05.

**Supplementary** Figure 9: (A) Representative immunofluorescent images corresponding to *in vitro* adult mouse CM proliferation assay. EdU-positive nuclei are marked as red, the CMs were visualized through Troponin I staining (green), and all the nuclei were stained by DAPI (blue). (B) The dot plot represents the quantitative analysis of EdU-positive CM in *MCM2* overexpressing CM versus control. (C) Representative immunofluorescent images corresponding *in vitro* neonatal rat ventricular CM proliferation assay. KI67-positive nuclei are marked as red, the CMs were visualized through Troponin I staining (green), and all the nuclei were stained by DAPI (blue). (D) The dot plot represents the quantification of KI67-positive CM in *MCM2* overexpressing neonatal rat ventricular CM versus control. Data represented as mean±SE. * = p-value ≤0.05. # = statically nonsignificant. Statistical significance was calculated through the *T*-test. *P* value ≤ 0.05 was considered statistically significant. Scale bars represent the magnification of the corresponding image. All the experiments were performed in duplicate and repeated at least three times.

**Supplementary** Figure 10: (A-C) Dot plots representing the different parameters of tube formation analysis. More than 10 images per group were analyzed for vascular density analysis. Data represented as mean±SE. N ≥ 10 images per group. Statistical significance was calculated through a T-test to compare the data between the groups. P value ≤ 0.05 was considered statistically significant. * = p-value ≤0.05.

**Supplementary** Figure 11: (A) Representative immunofluorescent images, corresponding to *in vitro* proliferation assay after adding the conditioned media from the *MCM2* overexpressing adult mouse CMs (*MCM2*-CoMed) on endothelial cells (HUVEC). KI67-positive nuclei are marked as red, whereas, endothelial cells are visualized through vimentin staining (green), and all the nuclei are stained by DAPI (blue). (B) The graph represents the quantification of KI67-positive HUVEC after *MCM2*-CoMed treatment *versus* control. Data represented as mean±SE. Statistical significance was calculated through the *T*-test. *P* value ≤ 0.05 was considered statistically significant. # = statically nonsignificant. Scale bars represent the magnification of the corresponding image.

**Supplementary** Figure 12: Bubble plot demonstrating the cellular and sub-cellular association of MCM2 interacting proteins, which was analyzed through cellular component analysis. The size of the bubble corresponds to the number of genes in that component. Whereas, the color of the bubble corresponds to the significance value.

