## Supplementary Figures for "*MCM2* mediates post-MI cardioprotection by promoting the pro-angiogenic cardiosome signaling"

### Figure S1

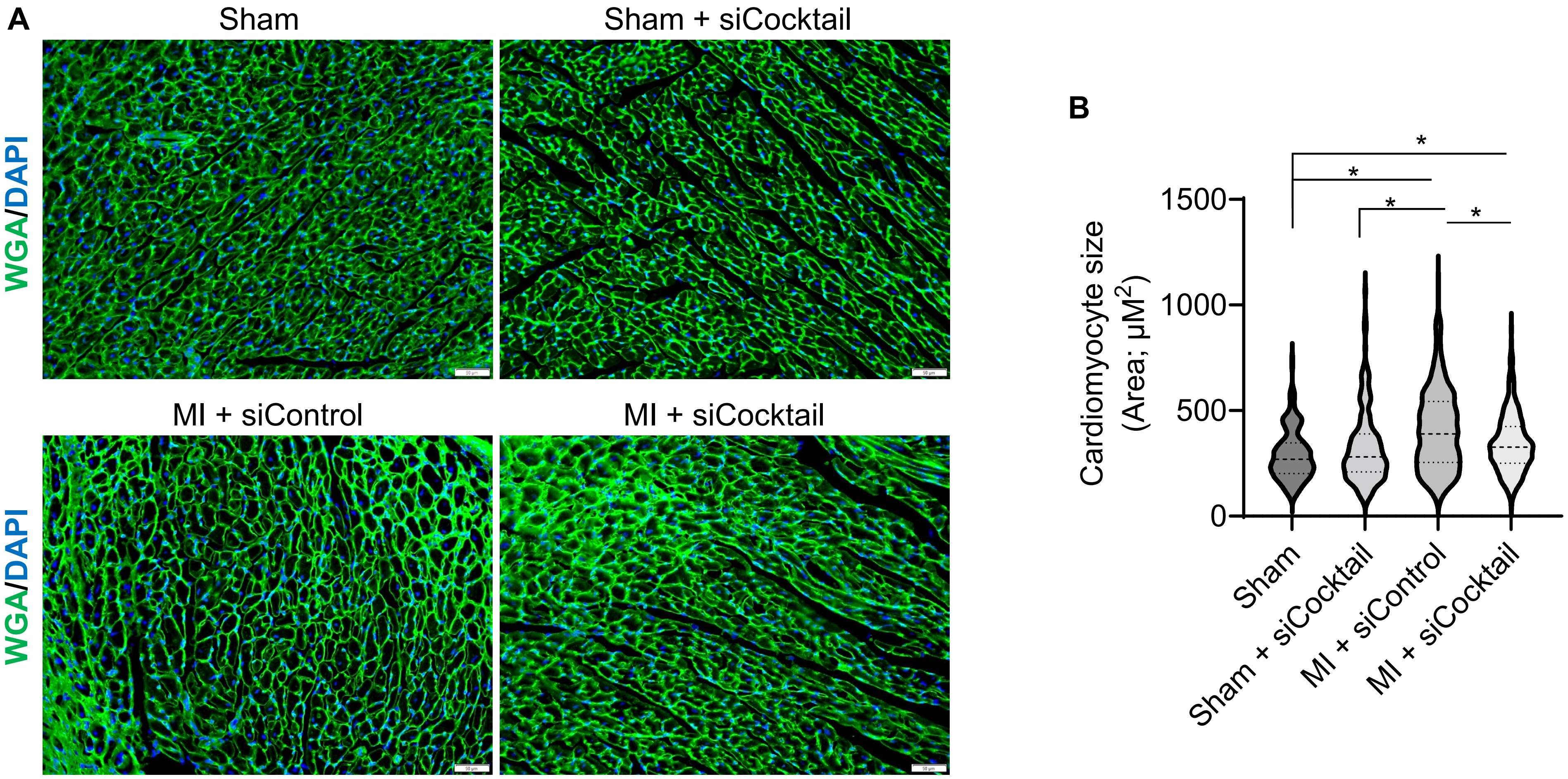
A
Sham
Sham + siCocktail
WGA/DAPI
MI + siControl
MI + siCocktail
WGA/DAPI
B
Cardiomyocyte size
(Area; μM2)
1500
1000
500
0
*
*
*
*
Sham
Sham + siCocktail
MI + siControl
MI + siCocktail

### Figure S2

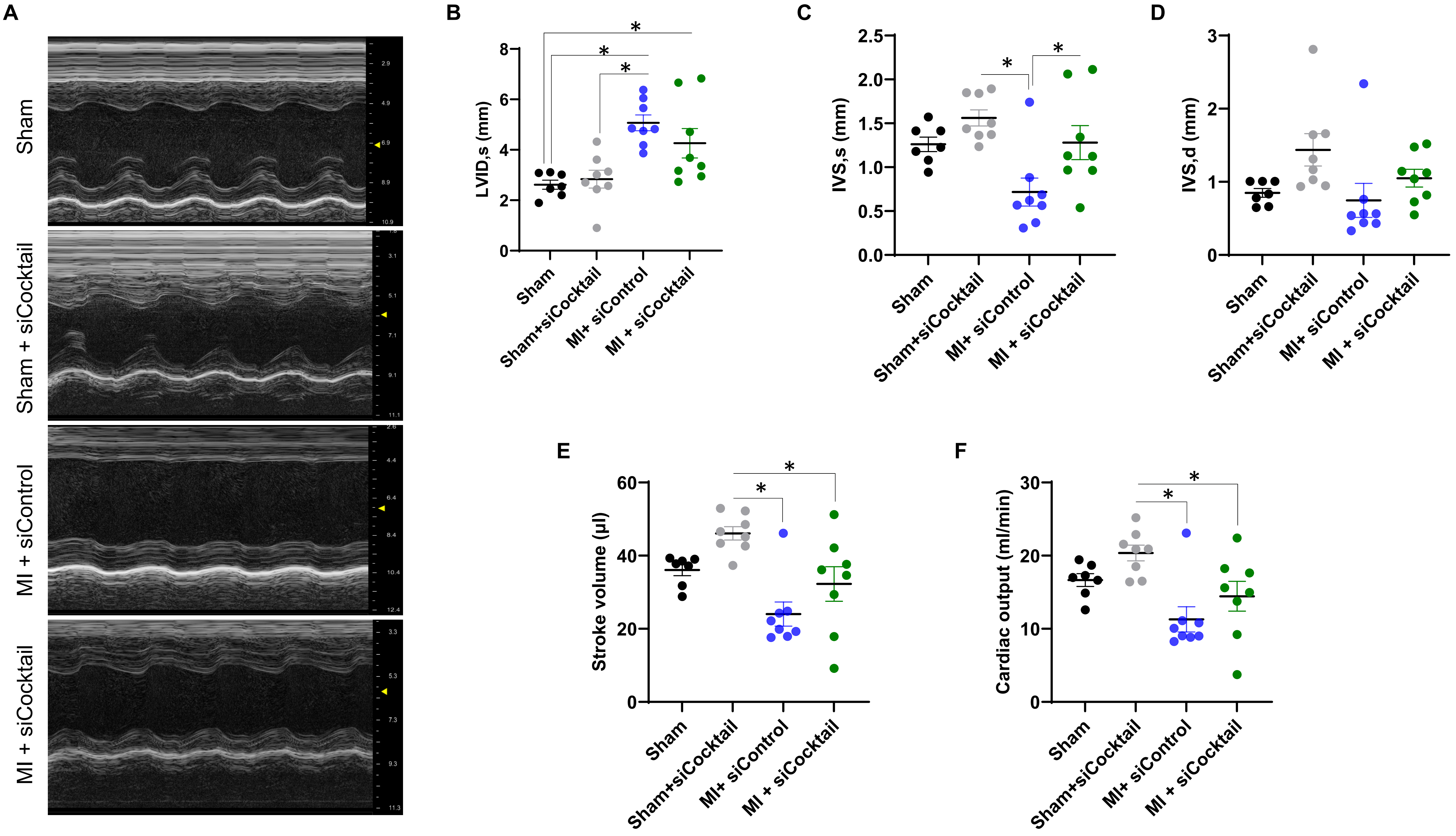

A
Sham
Sham + siCocktail
MI + siControl
MI + siCocktail
B
LVID,s (mm)
C
IVS,s (mm)
D
IVS,d (mm)
E
Stroke volume (µl)
F
Cardiac output (ml/min)
Sham
Sham+siCocktail
MI+ siControl
MI + siCocktail
*

### Figure S3

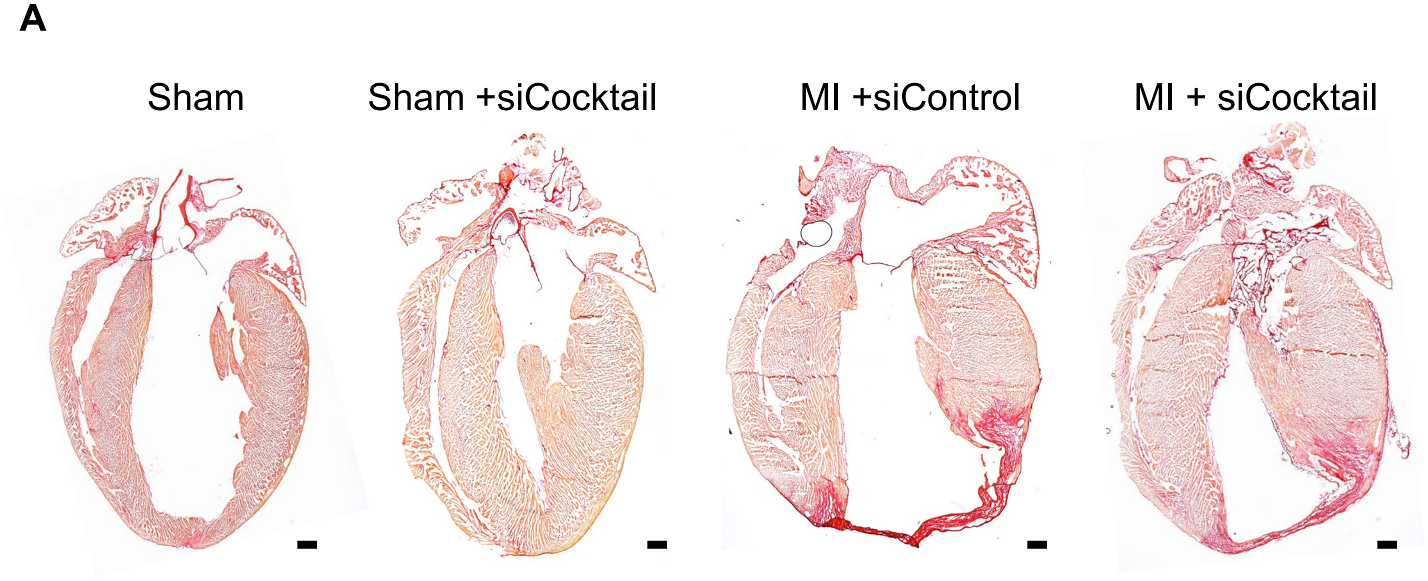
A
Sham
Sham +siCocktail
MI +siControl
MI + siCocktail

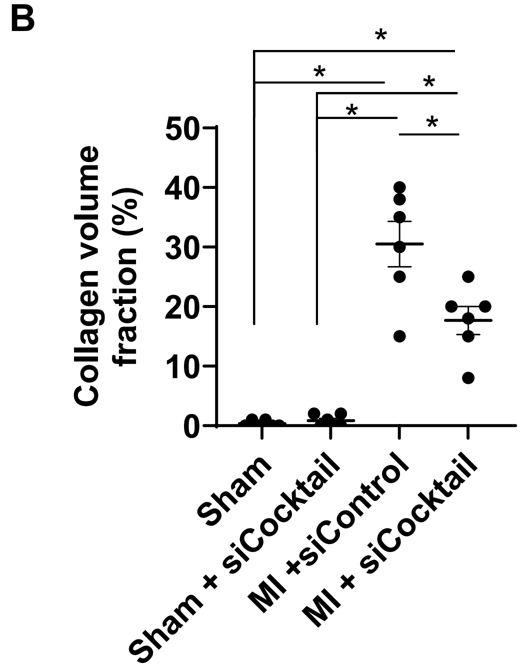
B
Collagen volume fraction (%)
50
40
30
20
10
0
*
*
*
*
*
Sham
Sham + siCocktail
MI +siControl
MI + siCocktail

### Figure S4

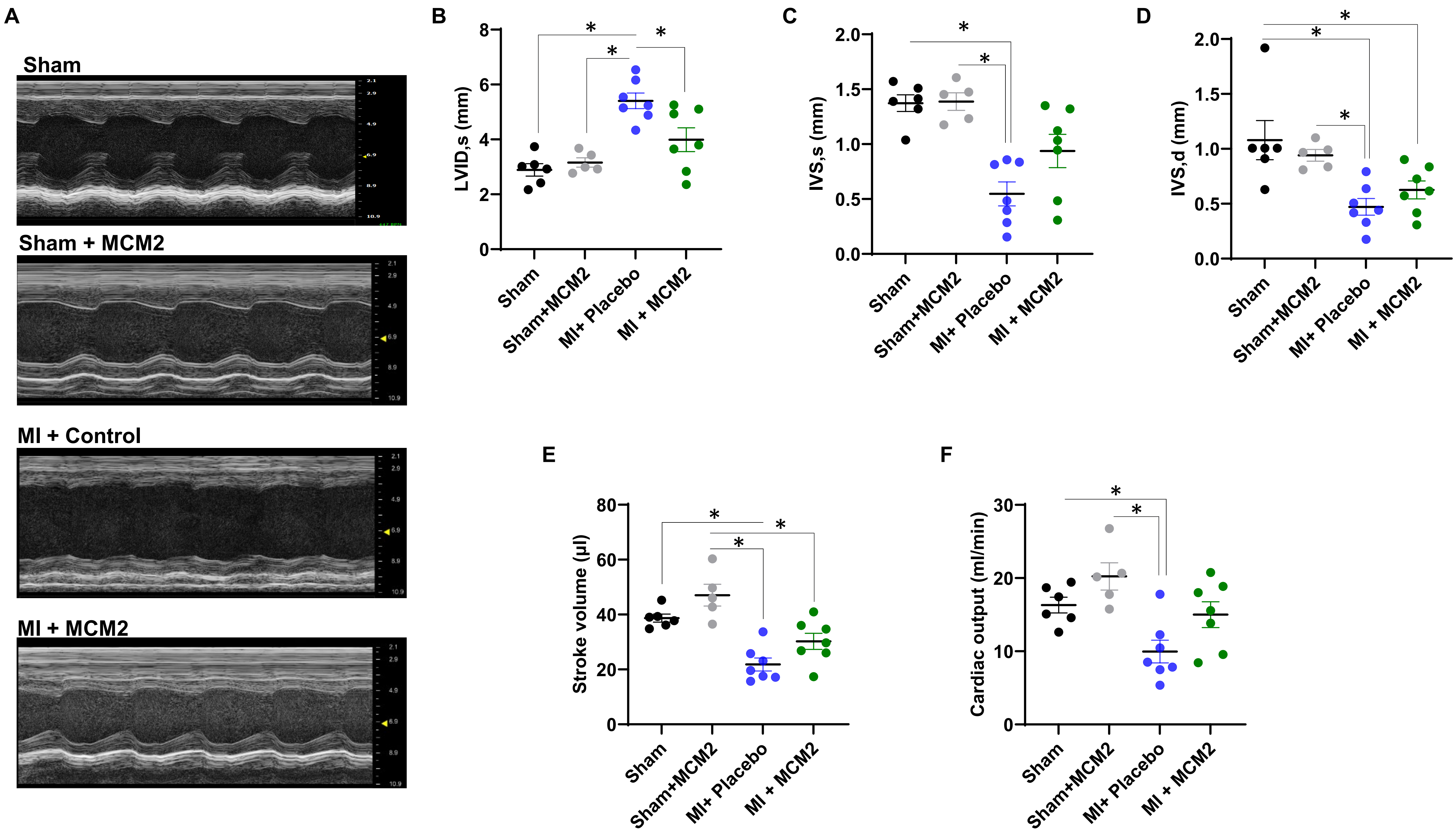
A
Sham
Sham + MCM2
MI + Control
MI + MCM2
B
LVID,s (mm)
Sham
Sham+MCM2
MI+ Placebo
MI + MCM2
C
IVS,s (mm)
D
IVS,d (mm)
E
Stroke volume (µl)
F
Cardiac output (ml/min)

### Figure S5 Overexpression of *MCM2* improves cardiac wall function after MI

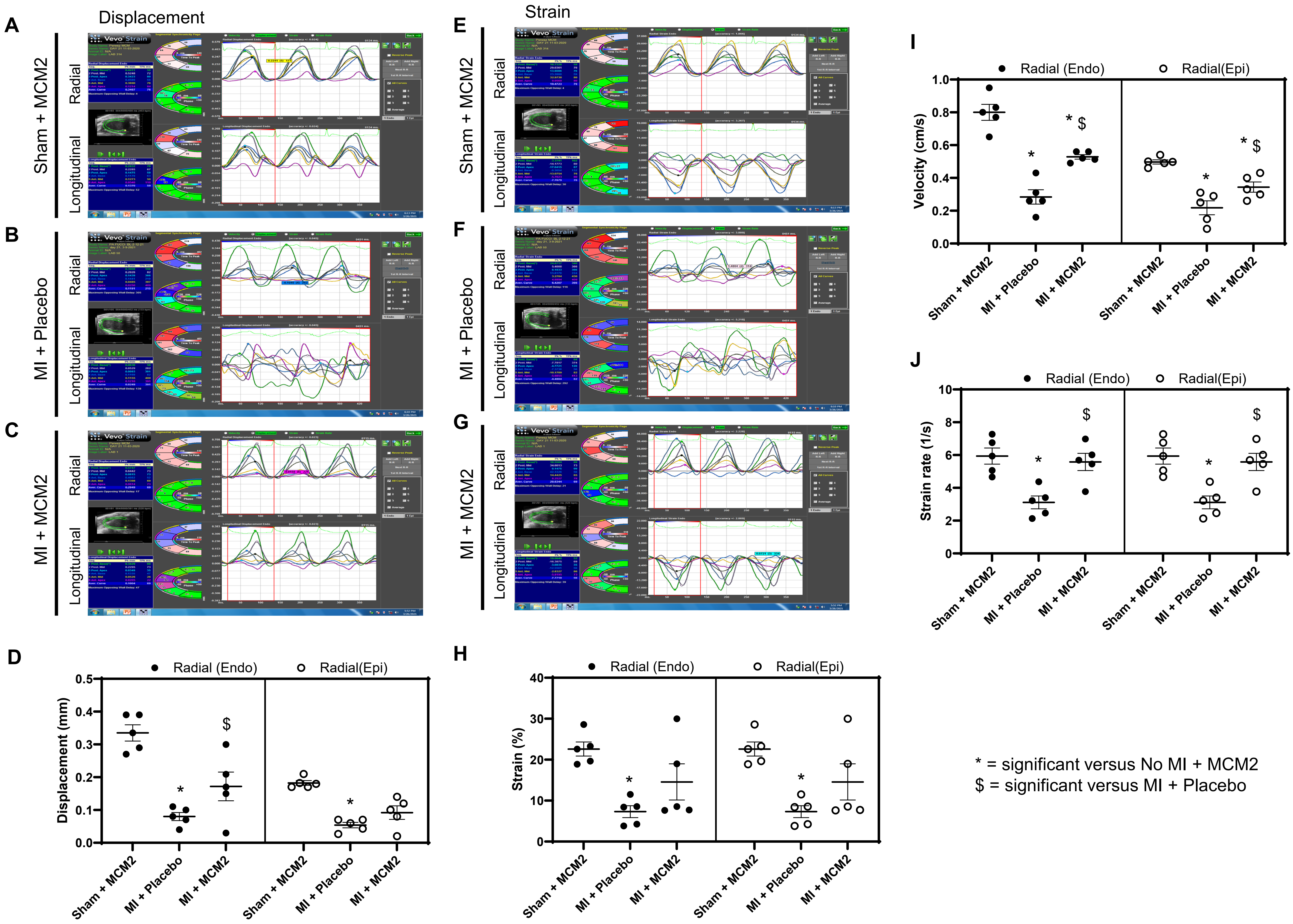

Figure S6

Overexpression of *MCM2* reduce cardiac remodeling in injured heart

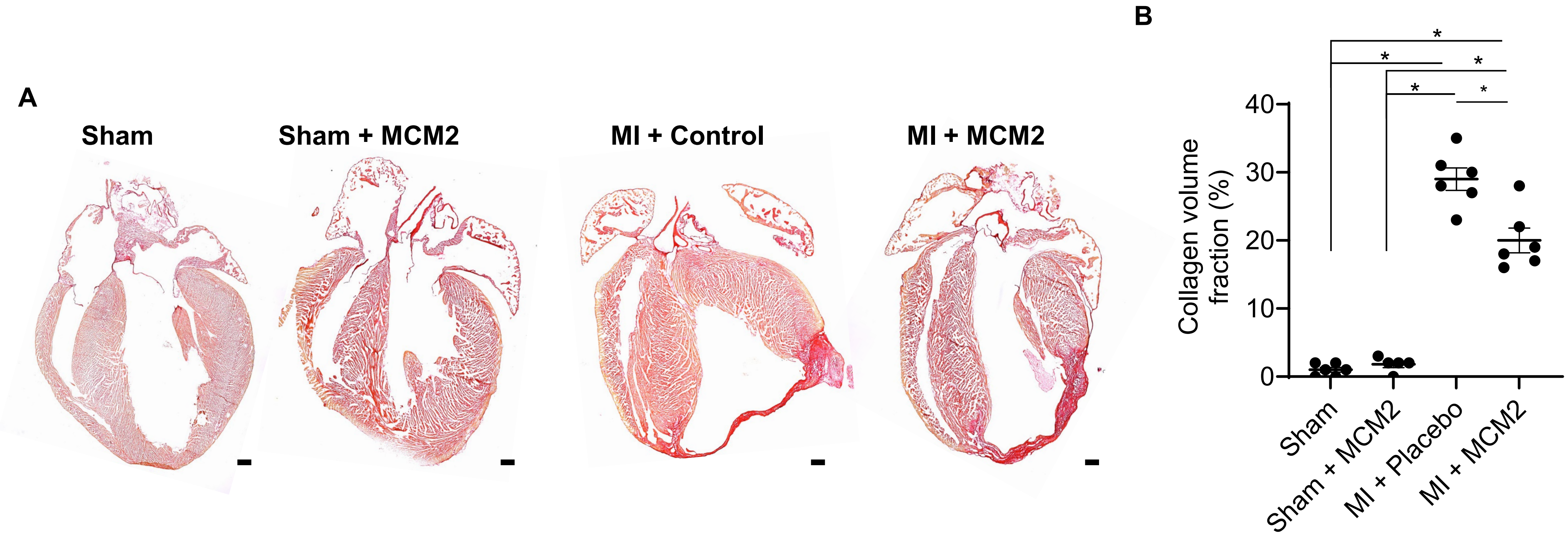

### Figure S7

#### Overexpression of *MCM2* is inadequate to induce cardiomyocyte proliferation: *in vivo*

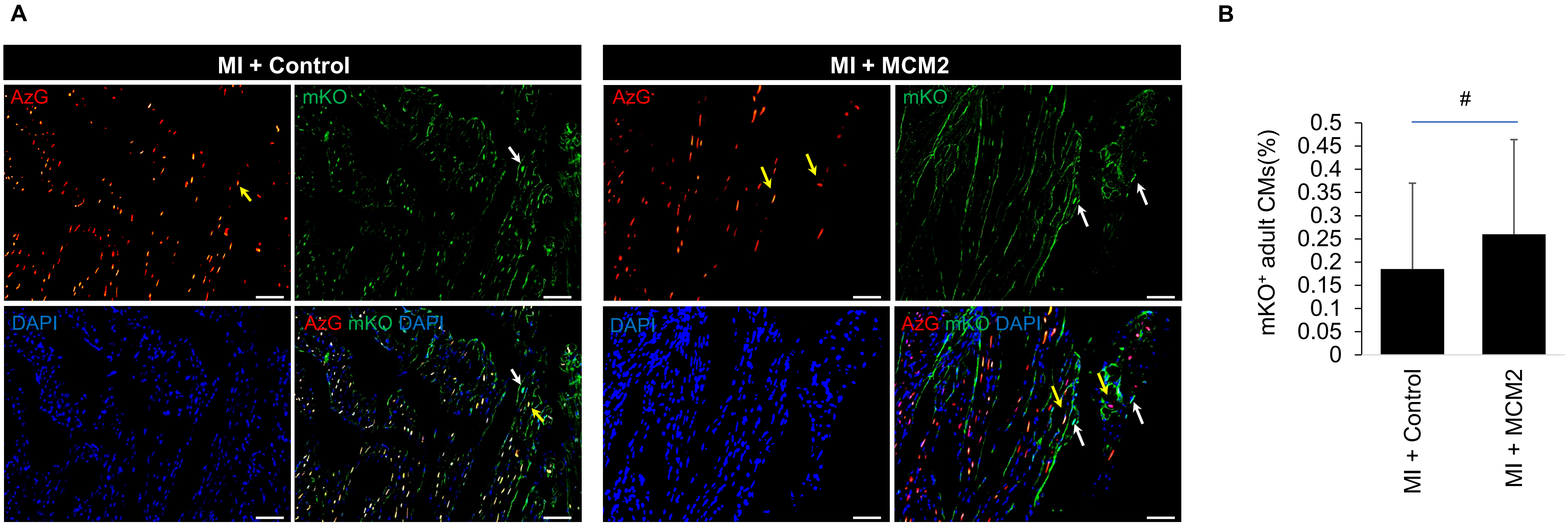

### Figure S8

**Overexpression of *MCM2* is inadequate to induce cardiomyocyte proliferation: *in vitro***

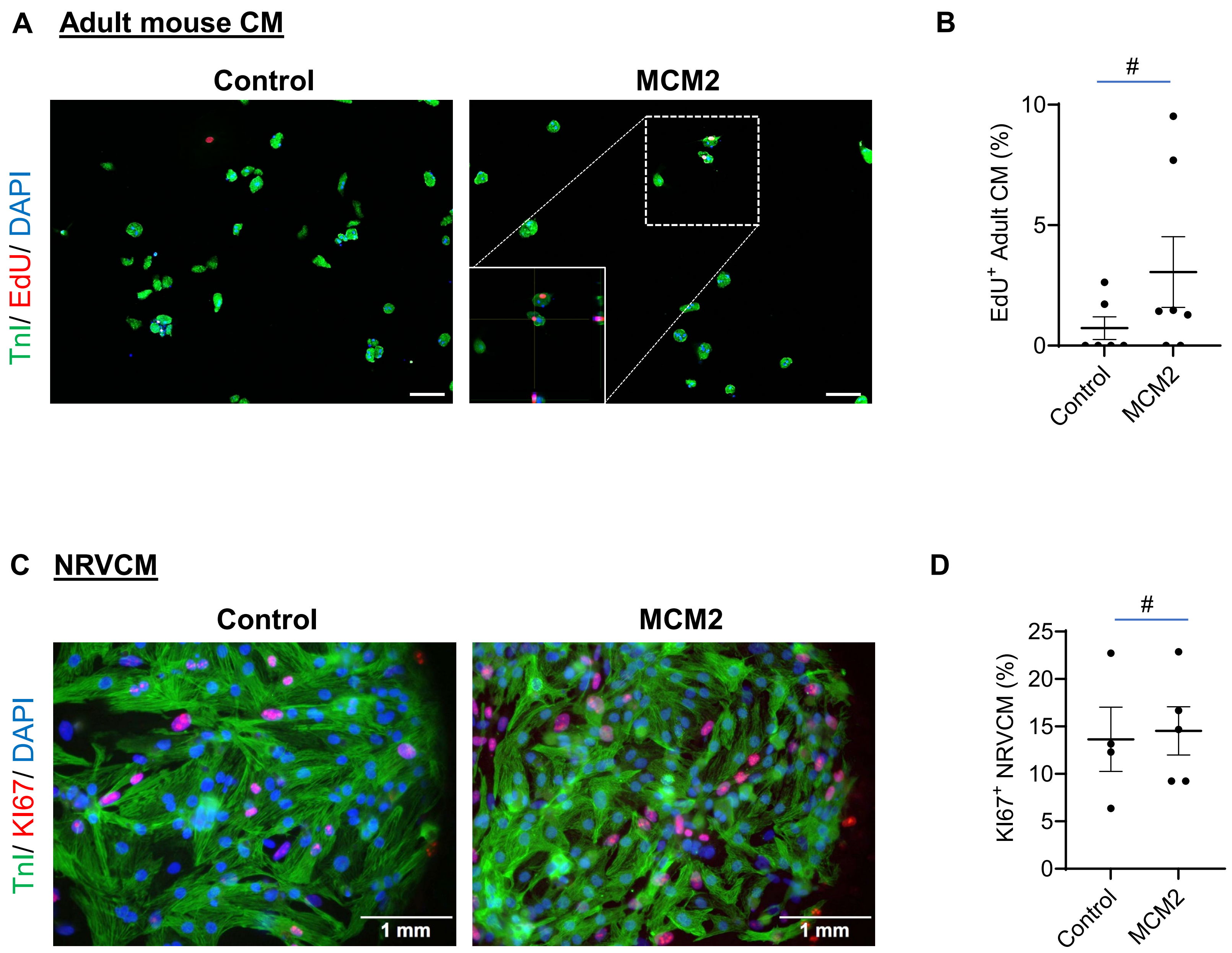

### Figure S9

#### *MCM2* overexpression in adult cardiomyocytes overexpression improves angiogenesis

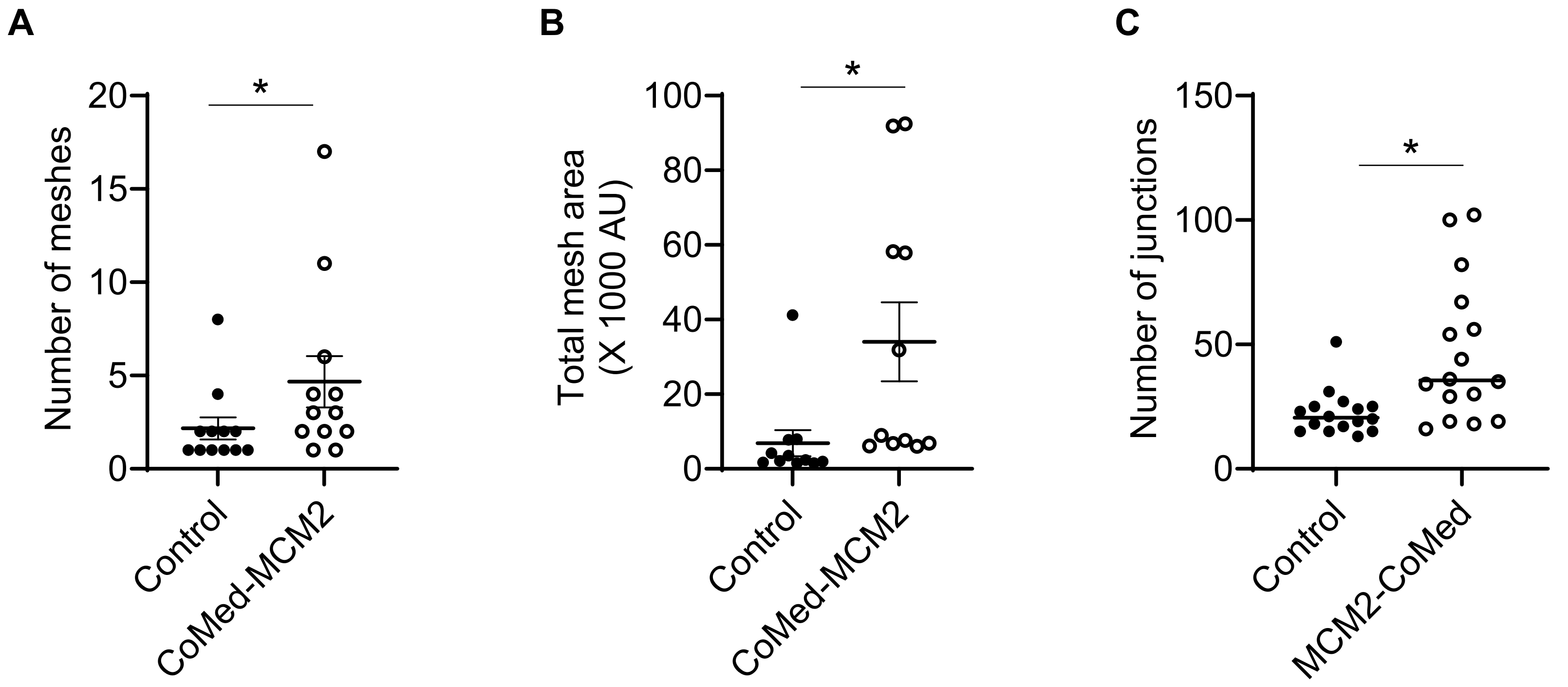


#### Conditioned medium from MCM2 overexpressing CM, is inadequate to increase endothelial cell proliferation

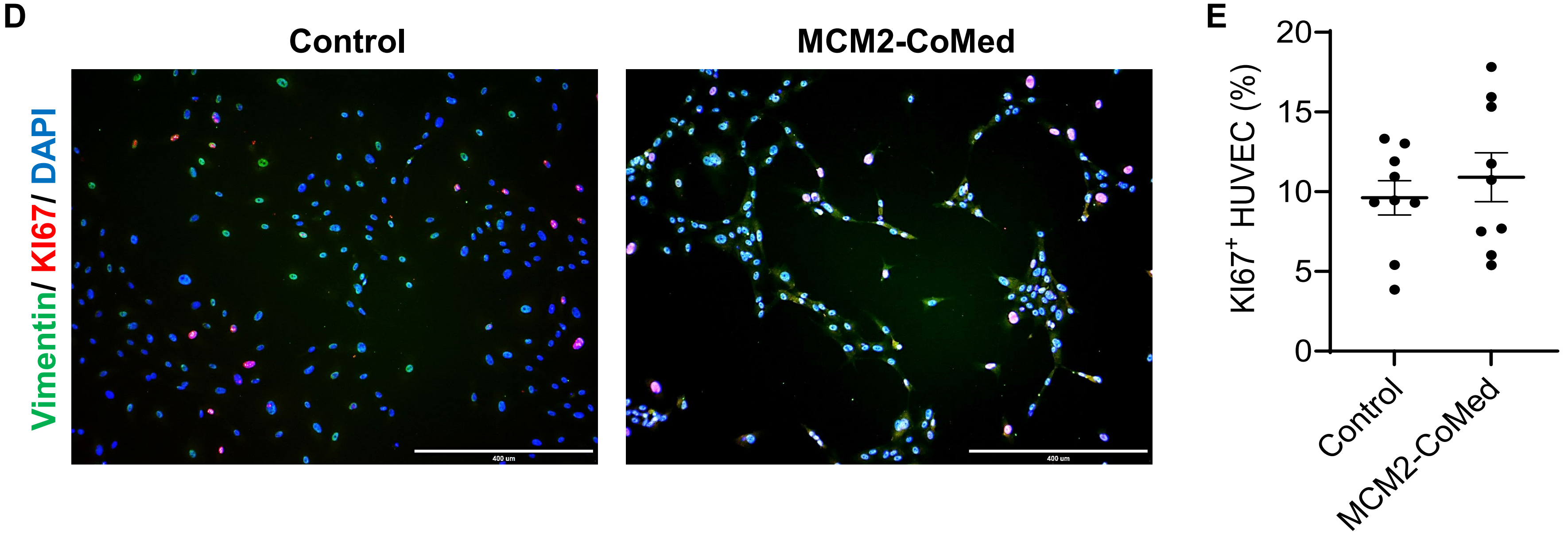

Figure S10

Cellular component analysis of MCM2 interacting proteins

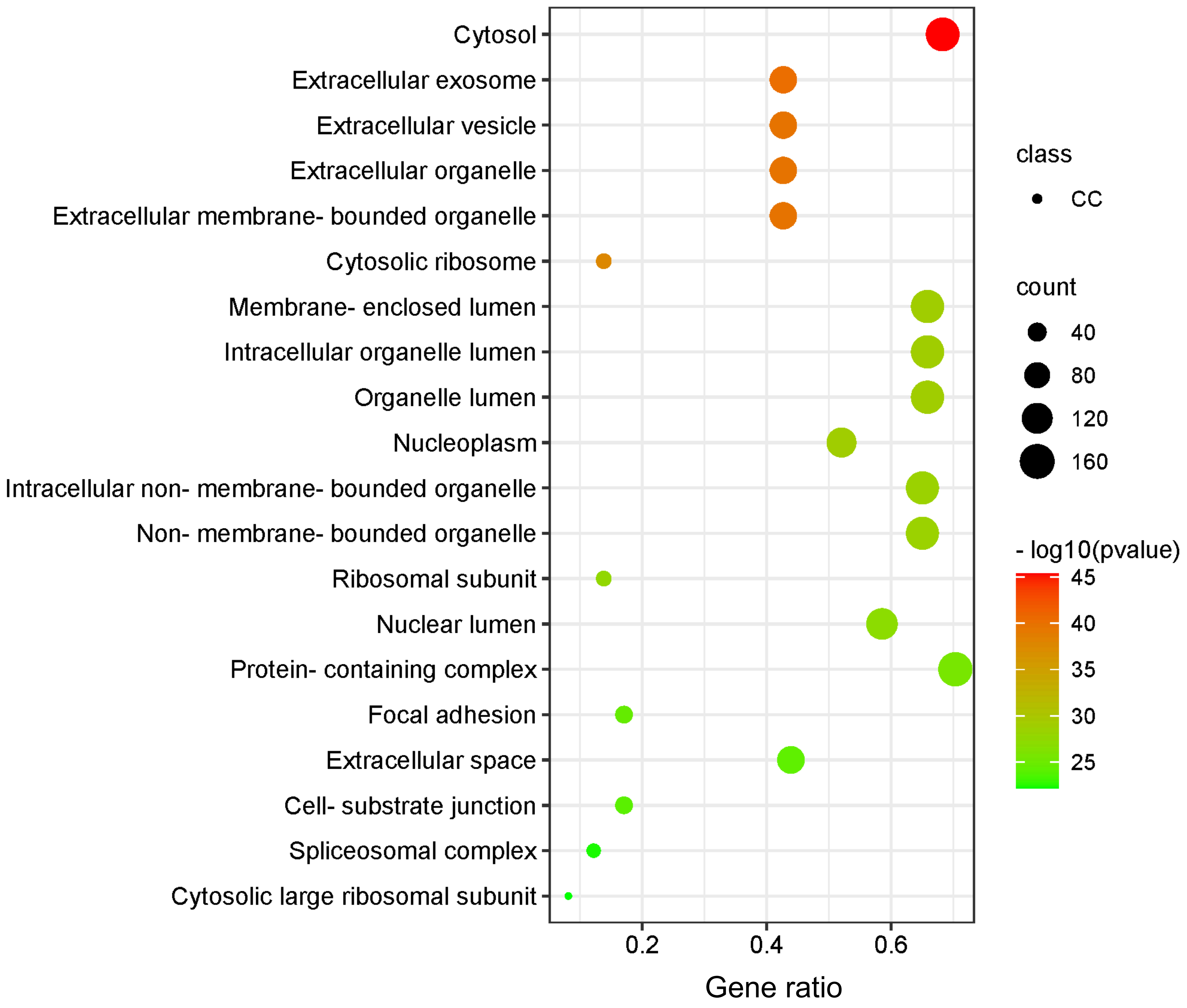
